# Scalable Screening of Ternary-Code DNA Methylation Dynamics Associated with Human Traits

**DOI:** 10.1101/2024.05.17.594606

**Authors:** David C Goldberg, Cameron Cloud, Sol Moe Lee, Bret Barnes, Steven Gruber, Elliot Kim, Anita Pottekat, Maximillian S Westphal, Luana McAuliffe, Elisa Majounie, Manesh KalayilManian, Qingdi Zhu, Christine Tran, Mark Hansen, Jelena Stojakovic, Jared B Parker, Rahul M Kohli, Rishi Porecha, Nicole Renke, Wanding Zhou

## Abstract

Epigenome-wide association studies (EWAS) are transforming our understanding of the interplay between epigenetics and complex human traits and phenotypes. We introduce the Methylation Screening Array (MSA), a new iteration of the Infinium technology for scalable and quantitative screening of trait associations of nuanced ternary-code cytosine modifications in larger, more inclusive, and stratified human populations. MSA integrates EWAS, single-cell, and cell-type-resolved methylome profiles, covering diverse human traits and diseases. Our first MSA applications yield multiple biological insights: we revealed a previously unappreciated role of 5-hydroxymethylcytosine (5hmC) in trait associations and epigenetic clocks. We demonstrated that 5hmCs complement 5-methylcytosines (5mCs) in defining tissues and cells’ epigenetic identities. In-depth analyses highlighted the cell type context of EWAS and GWAS hits. Using this platform, we conducted a comprehensive human 5hmC aging EWAS, discovering tissue-invariant and tissue-specific aging dynamics, including distinct tissue-specific rates of mitotic hyper- and hypomethylation rates. These findings chart a landscape of the complex interplay of the two forms of cytosine modifications in diverse human tissues and their roles in health and disease.

## INTRODUCTION

The dynamic genome-wide patterns of cytosine modifications, including 5-methylcytosine (5mC), 5-hydroxymethylcytosine (5hmC), and unmodified cytosine (collectively referred to as the ternary code methylation), play a critical role in regulating gene expression regulation^1^, genome stability maintenance^2^, and organismal development^3^. Through these roles, DNA methylation has been extensively associated with cellular and physiological human traits^4^ and is increasingly utilized as a biomarker in translational research and clinical applications^5,6^. Notable examples include applying DNA methylation to classify cancer and rare diseases^7–10^, liquid biopsy-based disease diagnosis^11^, and assessing disease hazard through methylation risk scores^12^ and forensic analysis^13^. Analysis of DNA methylation profiles is also crucial for elucidating gene transcription mechanisms^14^, understanding cell identity maintenance^15^, studying variations in cell composition^16^, and investigating gene-environment interactions within populations^4^.

Epigenome-wide association studies (EWAS) investigate large human populations for how DNA cytosine modifications are associated with human traits and diseases^4,17,18^. Over the past decade, EWAS has been instrumental in uncovering links between DNA methylation and diverse human phenotypes. To support these studies, methodologies developed to profile DNA methylation across the genome^19^ are often challenged by the large size of the human genome, the complex methylation biology across genomic regions, and prevalent inter-cellular heterogeneity in tissues^20^. The most comprehensive DNA methylation profiling assay is single-cell whole genome methylation sequencing (scWGMS), which offers unparalleled detail by providing base-resolution data for individual cells^21^. However, the high costs and technical complexity of scWGMS often limit its use to a limited number of samples^22^. As it is currently not practical to implement scWGMS for population studies, alternative methodologies are more frequently used, trading off genome coverage, base resolution, or cell-type resolution to reduce costs and technical demands. These include methods for profiling bulk tissues^23^ or FACS-purified cells (*e.g.,* bulk deep WGBS or nanopore sequencing)^24^, targeted genome capture (*e.g.,* RRBS^25^), and the use of data techniques to interpret sparse signals (*e.g.,* low-pass sequencing^26^).

The Infinium DNA methylation BeadChip has been a robust solution for large-scale methylation discovery and screening efforts due to its ease of experiment and data analysis^27^, base-resolution detection, and high quantitative granularity. This platform has been central to consortia such as The Cancer Genome Atlas (TCGA) and has amassed over 80,000 HM450 methylomes^28^ and a comparable number of EPIC array methylation profiles in the Gene Expression Omnibus (GEO). While sequencing-based methods are more used for case-specific and mechanistic studies, Infinium arrays are often preferred in discovering population-scale trait associations, including meQTL studies^29,30^, epigenetic risk scoring^31,32^, and EWASs in human^33,34^ and other mammalian species^35–37^. Such adoption is partly due to the need for population studies to cover a large number of samples to dissect multiple cohort covariates (*e.g.*, sex, age, genetic background, and tissue type) and their interactions and, in others, to the high depths required to capture nuanced variations in cytosine modification levels^38,39^. A prominent example is 5hmCs, which are inherently stochastic—often under 30% per site, even in homogeneous cell populations^40^, unlike the bimodal distribution typical of 5mCs—and are concentrated in specific regulatory regions^41,42^, necessitating high quantitative resolution for accurate measurements on a small number of sites rather than sparse whole-genome coverage.

Array technologies rely on static probe designs that fix the CpG space to those selected during the array’s development^43^. While this permits cross-study comparisons, the current design has the following limitations. First, whole genome methylation sequencing of 5mCs and 5hmCs in human cells and tissues has significantly advanced our understanding of cell type methylation at high resolutions^24^ since the last human array design^44^. Current EPICv2 arrays, largely inheriting EPIC, have yet to incorporate the recent discoveries (e.g., of 5hmCs)^22,24,38,39^. Further, most predictive models based on existing arrays hinge on a small number of trait associates. For example, most epigenetic clock models used hundreds of CpGs and reached high prediction accuracy^45^. Minimalistic approaches were taken in epigenetic clock construction^46^, cell type deconvolution^47^, and cancer classification^48^. These observations prompt us to think that building compatible but condensed arrays for applying existing models and reassessing associations in significantly larger, more inclusive, and stratified human populations should be feasible (Figure 1A).

**Figure 1:**
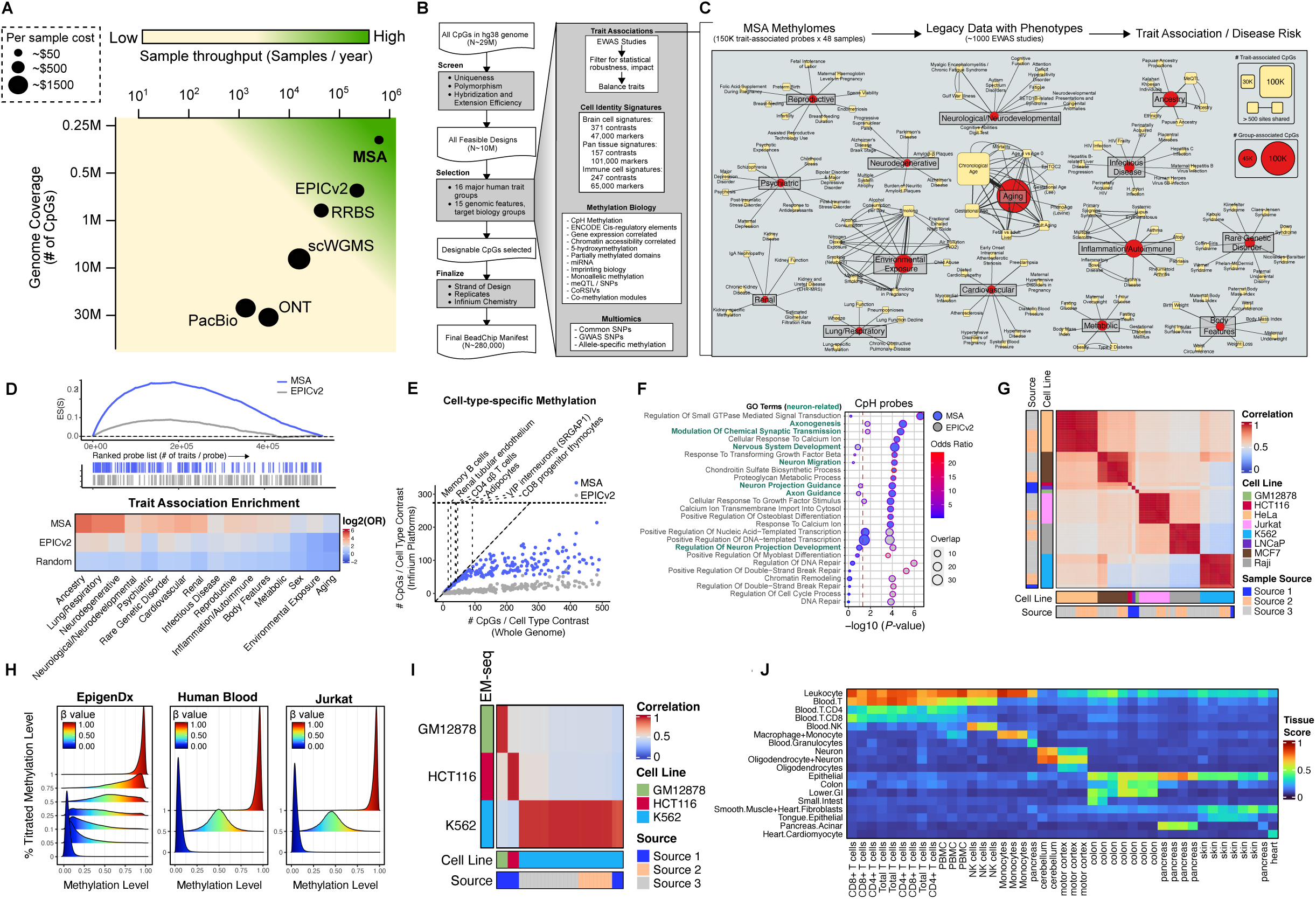
MSA design workflow and major trait groups. (A) Schematic illustrating the axes of sample throughput, genome coverage, and cost efficiency for different methylation assay technologies. MSA has targeted genome coverage for high throughput at a relatively cheaper cost. (B) The screening process is to identify designable probes (left), targeted trait groups, and methylation features (right). (C) Major trait categories are incorporated into array content with a subset of represented sub-trait groups. Two trait categories (Sex and Other) are omitted. Duplicate traits studied from different cohorts are possible. (D) Set enrichment analysis showing the enrichment score of MSA and EPICv2 probes down a ranked list of EWAS hits probes, ranked according to the number of trait associations (top) and heatmap showing the enrichment of retained sites on MSA in all annotated major trait groups compared with EPICv2 and a random selection of Infinium probes equal in size to MSA (bottom). (E) Number of CpGs per cell type contrast on MSA vs. EPICv2 for contrasts with few (<500) high-quality whole genome markers. (F) Gene ontology for biological process results for genes linked to CpH probes (minimum two probes per gene) on MSA and EPICv2. (G) Heatmap of beta value correlations between cell line samples profiled on MSA. (H) Density plots of beta values for methylation titration standards. (I) Heatmap of beta value correlations between cell line samples profiled on MSA and with an EM sequencing protocol. (J) Tissue scores for a subset of profiled tissues were generated using the nearest neighbor probes on MSA for an EPIC tissue prediction model.

To implement these thoughts, we present the rational, systematic design and the first application of the Methylation Screening Array (MSA), the latest Infinium BeadChip iteration. Compared to previous Infinium BeadChips, MSA has concentrated its coverage on trait-associated methylation (∼5.6 trait associations per site vs. ∼2.2 in EPICv2, Methods) and cell-identity-associated methylation variations (∼3.7 cell signatures per site vs. ∼2.3 in EPICv2, with an additional 48 novel cell type contrasts that can be made). Half of the design targeted previously reported EWAS associations. The other half leverages the latest single-cell and bulk whole genome methylation profiling efforts that deeply characterize diverse human cell types. This dual approach enables high-resolution cell-type deconvolution, supported by reference methylation panels and predictive models we have rigorously benchmarked in this study. Compared to the 8-sample plate design used in previous methylation arrays, MSA is built on a novel 48-sample EX methylation platform to achieve ultra-high sample throughput at a lower cost per sample while screening for more traits per probe site. Evaluation of the array’s accuracy and reproducibility confirms its robustness for population-scale applications. Applying MSA to various human tissues, we characterize tissue-specific 5mC and 5hmC genomic distribution and demonstrate the capacity for accurate cell-type deconvolution. We performed the first EWAS for 5hmC in aging and sex and identified previously under-reported contributions of 5hmC to the prediction mechanism of epigenetic clocks. Analysis of 64 whole blood methylomes demonstrated variable methylation at established EWAS loci and age and sex-related immune cell composition alterations across the lifespan.

## RESULTS

### Rational, systematic design of MSA

We designed the MSA array by compactly consolidating human trait-associated loci identified in previous EWAS studies and novel probe designs targeting diverse methylation biology (Figure 1B). After post-manufacture quality control, the MSA array contains 284,317 unique probe sets targeting 269,094 genomic loci. 145,318 loci overlap what is targeted by the EPICv2 platform (Figure S1A). More SNP-targeting probe sets and nearly as many CpH probes were incorporated relative to EPICv2 (Figure S1B). Human trait-associated methylations were identified by mining EWAS databases and literature, prioritizing the diversity of trait coverage and statistical significance (Methods). We broadly classified all EWAS hits into 16 trait groups (Figure 1C, S1C). As expected from the design, MSA is highly enriched by EWAS associations across human traits (Figure 1D), reflecting the platform’s targeted design and compact size.

For new CpGs that previous Infinium platforms have not targeted, we leveraged existing WGBS data sets to identify CpGs associated with cell type, cis-regulatory elements, correlation with chromatin accessibility and gene expression, 5-hydroxymethylation and additional methylation features (Figure S1D, Methods). We emphasized high-confidence cell-type-specific methylation discriminants to facilitate the deconvolution of complex heterogeneous tissue types and the study of cell-specific processes. Using pseudo bulk and sorted methylomes from brain^49–51^, pan tissue^24^, and blood cells^52^, we performed hierarchical, non-parametric analyses to identify CpG discriminants for the different cell types (Methods). These analyses identified thousands of hyper and hypomethylated signatures across hundreds of cell types (Figure S1E). Compared to EPICv2, MSA contains more markers per cell type comparison group despite the smaller size (Figure 1E). These differences are especially pronounced for rarer cell types or comparison groups with relatively few designable genome-wide markers. For example, our analysis of WGBS data identified 34 high-quality markers of the SRGAP1 subtype of VIP interneurons derived from the caudal ganglionic eminence. We incorporated 31 markers onto MSA, whereas EPICv2 contains three (Figure 1E).

Like the EPICv2 array, the MSA design is highly enriched in the promoter, enhancer, and transcriptionally active regions. It is strongly depleted from quiescent, heterochromatic, and ZNF regions (as annotated by the full stack ChromHMM^53^) (Figure S1F, Table S1). The two platforms are less represented by open-sea CGI sites but have a higher proportion of cis-regulatory element coverage (as annotated by ENCODE^54^) (Figure S1G, Table S1). MSA has a slightly increased proportion of proximal (5.6% vs. 3.45%) and distal (16.2% vs. 10.1%) enhancer elements and marginally less coverage of CpG island (12.4% vs. 16.2%) sites compared to EPICv2. Compared to EPICv2, MSA CpH probes were designed by analyzing brain cell type-specific methylomes with more prevalent CpH methylation. The queried cytosines are more linked to brain and neuron functions, implicating genes critical for neuron development and synaptic signaling (Figure 1F).

Lastly, MSA contains at least one probe linked to 14,964 genes (overlapping or within 1500bp of the transcription start site), nearly as many as the larger EPICv2 array (Figure S1H). The 772 genes on EPICv2 but not MSA were enriched in olfactory receptors and highly polymorphic genes whose readings are often confounded by genetic polymorphism^55^ (Figure S1I). In summary, the MSA assay targets human trait-associated methylations and novel sites where methylation is predicted to be dynamic, cell type-specific, and biologically relevant.

### MSA is highly reproducible and accurate

We used the MSA BeadChip to generate 146 methylation profiles for eight cell lines (GM12878, HCT116, HeLa, Jurkat, K562, LNCaP, MCF7, and Raji) with replicates in order to assess the arrays’ technical performance. Probe success rates for most of these 146 samples surpassed 90% (Methods, Figure S1J). Probe detection rates were robust to 50 ng of input DNA but declined to <60% for three samples with ∼30 ng of input DNA (Figure S1J).

For all cell lines, we observed high correlation coefficients between samples of the same line regardless of the laboratory of cell culture (Figure 1G). The correlations between different cell lines were significantly lower, reflecting the different cell origin, ploidy, and epigenomic properties of the different lines. For GM12878 and HCT116, we generated technical replicate methylomes using the same DNA sample and computed Spearman correlation coefficients and F1 scores based on binarized methylation levels (Methods). The technical replicates had highly similar methylation profiles, with Spearman’s rho of 0.986 and 0.945 and F1 scores of 0.99 and 0.95 for GM12878 and HCT116, respectively (Figure S1K). We also compared the GM12878 cell line to methylation profiles that we previously generated using the same DNA samples on the EPIC and EPICv2 array^44^. The technical replicate correlation coefficients surpassed 97% on all three platforms. Across shared probes, methylation measurements were highly concordant between platforms (Figure S1L).

Like the EPICv2 BeadChip, the MSA array includes replicate probe designs that target the same 122-mer genomic loci but may vary in the other design details^44^. The replicate designs have the same prefix but alternative suffixes that describe the chemistry and target strand specifications^55^. For each of the 8,523 replicate probe groups, we calculated the standard deviation (SD) of replicate probes within cell line samples and compared the means of these SDs to the SDs of non-replicate probes (Figure S1M). Replicate probes had a low mean standard deviation of 0.02 compared to non-replicate probes, suggesting that the replicate probes produce consistent methylation measurements. Methylation can be averaged over replicate probes or the most robust replicate selected based on signal intensity *P*-value using *SeSAMe*^56^.

We also assessed the specificity of MSA probe sequences. To minimize cross-hybridization, only sequences with mapping quality >20 were considered for novel probe designs (Methods). In the final MSA manifest, >99.9% of probe sequences are uniquely mapped with high quality. The minority of probes with lower-quality mapping can be readily identified in the standard SeSAMe^56^ preprocessing pipeline.

Next, we evaluated the accuracy of MSA by comparing MSA beta values with methylation titration standards. For each titration, the beta value distributions centered on the target titration level (Figure 1H). We further compared our cell line methylomes from MSA to methylomes of the same DNA samples generated using an EM sequencing protocol^57,58^ (Figure 1I). Beta values remained highly correlated within but not across cell lines. Additional comparisons of the MSA cell line methylomes with publicly available WGBS data of the same cell lines also showed higher intra-cell line correlations than between cell lines (Figure S1N). These experiments confirm that MSA measurements are accurate and consistent with ground truth titrations and WGBS data.

While MSA is more scalable than prior platforms due to its smaller size, a substantial number of probes were not reintroduced (Figure S1A), which could hinder the implementation of methylation-based prediction models or the study of prior associations. First, we noted that the loss of probes minimally affected the performance of eight prior epigenetic clocks (Figure S1O). We also reason that missing EPIC probes can be imputed from MSA probes. We implemented a sparse nearest-neighbor graph approach on a deep WGBS data set of sorted human cells^24^ with high coverage across both platforms (Methods). Of the 714,492 non-retained probe sites, 471,145 had a nearest neighbor with a correlation >.5 across the WGBS methylomes. To evaluate whether EPIC-based models could retain compatibility with MSA methylation profiles, we trained a tissue prediction model using only legacy probes. We tested the prediction on MSA-profiled human tissue types. The beta value reading at the nearest neighbor MSA site was sufficient to predict the tissue type using the EPIC-only model (Figure 1J). We have provided a neighbor reference in Supplementary Table S3 for missing value imputation.

### MSA uncovers tissue-specific methylation biology

We generated methylomes for five different sorted immune cell types (CD4 T, CD8 T, Total T cells, NK cells, Monocytes), peripheral blood mononuclear cells (PBMCs), and 26 different human tissue types (Figure S2A). We performed unsupervised clustering using t-stochastic neighbor embedding (tSNE) to explore their global methylome similarities. Related cell and tissue types were highly colocalized (Figure 2A).

**Figure 2:**
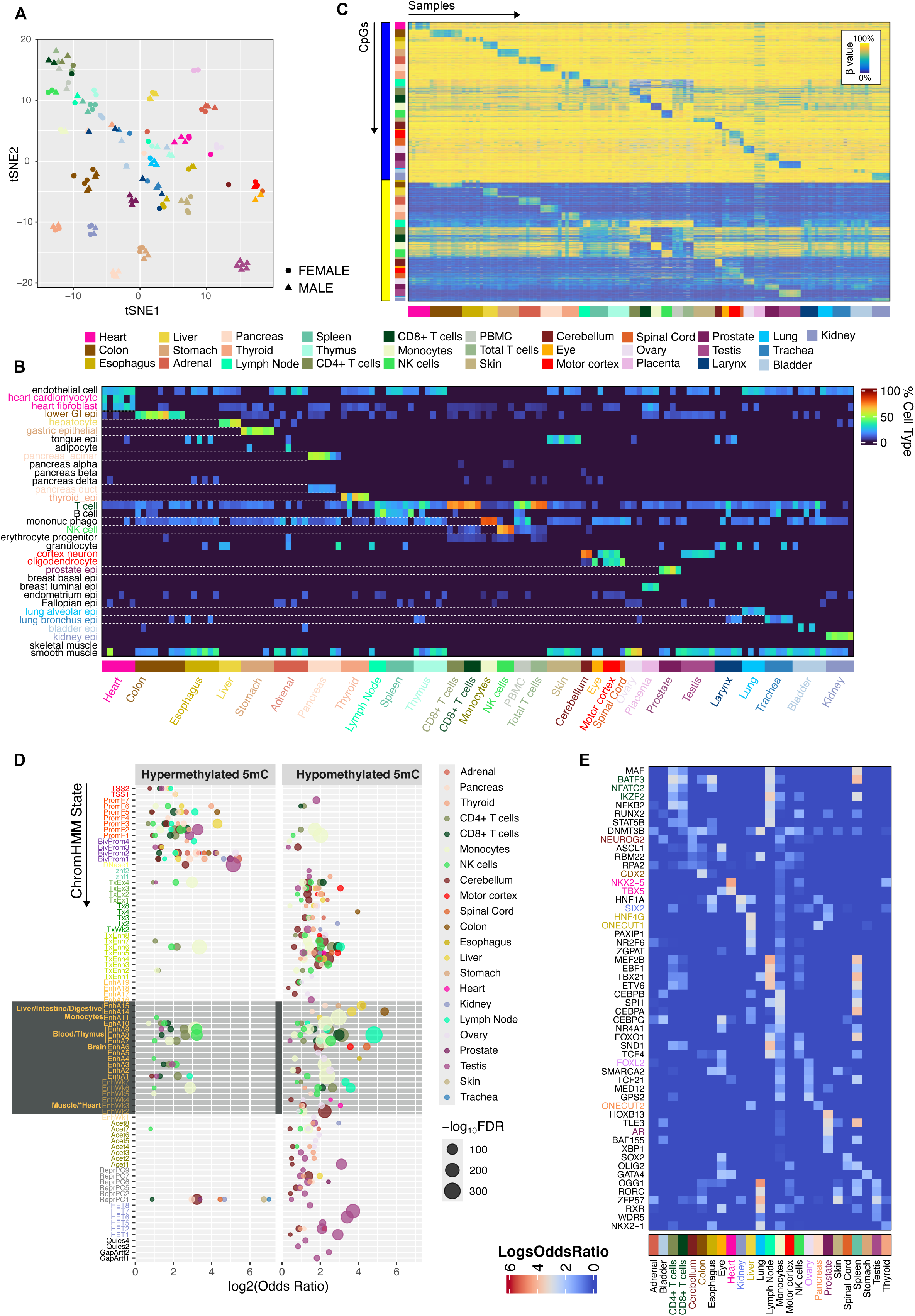
MSA reveals tissue-specific methylation biology and tissue compositions. (A) tSNE plot showing unsupervised clustering of sorted immune cells and bulk tissues profiled on MSA. (B) Heatmap showing cell type proportion estimates obtained by methylation-based deconvolution on the sorted immune cell and bulk tissues (columns) profiled on MSA (C) Heatmaps showing beta values of tissue-specific CpGs (rows) over bulk and sorted immune cells (columns). (D) Enrichment of hyper and hypomethylated tissue-specific CpGs in different full-stack ChromHMM chromatin states (FDR < .05) (E) Heatmap showing enrichment of tissue-specific hypomethylated CpGs (columns) in transcription factor binding sites (rows).

Cell type proportions are often the main drivers of bulk tissue EWAS results^59^. Using reference-based deconvolution, we tested whether our bulk MSA tissue methylomes could be resolved into their constituent cell types (Methods). The cell proportion estimates of bulk tissues aligned with the reported tissue types (Figure 2B). For example, heart samples were predicted to contain cardiomyocytes, heart fibroblasts, and endothelial cells, while liver samples primarily contained hepatocytes. As expected, organs of immune cell development, such as the spleen and lymph nodes, had varying proportions of monocytes, T cells, and B cells. The thymus lacked B cells, which is consistent with its role as an organ of T lymphocyte maturation^60^. A few profiled samples had discordant cell proportions and did not cluster in proximity to the rest of the samples of the same tissue type. For example, while most pancreatic tissues were estimated as acinar and ductal cells, the most populous cell types of the organ^61^, one sample had a higher fraction of granulocytes, suggesting excessive blood contamination or sample mislabeling. Such cases were indicated and excluded from downstream tissue-specific analyses (Methods).

Next, we performed one-vs-all non-parametric supervised analyses of the tissues (Methods) and identified thousands of CpG discriminants uniquely methylated in the target tissue type (Figure 2C). Most CpG signatures were hypomethylated compared to the remaining tissues (Figure S2B). These tissue-specific probe sets were highly enriched in the cell-specific CpG signature lists curated from the analysis of publicly available single and sorted cell data sets during array design (Figure S2C, Methods), validating the design process and the performance of the selected probes in discriminating the target cell types.

To explore the role of tissue-specific methylation markers in the corresponding tissue biology, we analyzed the chromatin state distributions and gene linkages of the CpG sets. We first compared them with the full stack ChromHMM states, a universal genome annotation learned from over 1,000 data sets comprising diverse cell types^53^ (Figure 2D). Hypermethylated tissue signatures were generally absent from enhancers and were enriched in promoter and bivalent promoter states, while hypomethylated markers were enriched in enhancers and gene bodies. The signatures are strongly enriched in the chromatin state associated with the matching cell type. For example, cerebellum and motor cortex signatures are enriched in EnhA6, representing brain enhancers. In contrast, colon and liver signatures were strongly enriched in EnhA14/A15, annotated as liver/digestive/intestine enhancers. The monocyte, NK cell, CD4+, and CD8+ T cell signatures were specifically enriched in EnhA7, a blood enhancer state.

In addition to tissue-specific chromatin states, the signatures colocalized with the corresponding tissue-specific transcription factor binding sites (Figure 2E). For example, CpG markers of kidney tissues were enriched in the binding sites of SIX2, which regulates the specification and maintenance of nephron progenitors^62^, while colon signatures were enriched in CDX2, which governs intestinal development and gene expression^63^. The markers were also in proximity to tissue-specific genes. We linked each tissue CpG marker to all genes within 10KB and co-embedded the linked gene sets with the human gene atlas ontology database (Figure S2D). Related tissue types are localized in the network space, and ontology terms match the tissue type. Collectively, our MSA data uncovered the epigenome signatures at tissue-specific transcription factor binding sites and genes that regulate the corresponding tissue biology.

Lastly, we analyzed the mitotic histories of the different tissue methylomes using a subset of PRC2 target CpGs^64^ and partially methylated domains (PMDs) to track the cumulative cell divisions of the tissue (Figure S2E). Applying the models to our tissue and immune cell methylomes yielded division rates consistent with the relative proliferative activity of these tissues reported in the literature based on radioisotope labeling^65^. For example, the colon, small intestine, and T cells had the highest division rate score, consistent with the high cellular turnover of these tissues (Figure S2E). In contrast, tissues with higher fractions of post-mitotic cell types, such as the motor cortex, cerebellum, and kidney, had the lowest division rates. The estimates of mitotic activity using the PMD methylations largely correlated with those obtained from the PRC2 model. Interestingly, pancreatic and adrenal tissues showed relatively low PMD methylation compared to other tissues and predictions based on average PRC2 target methylations. These effects were not fully explained by global methylation differences, which were minor for tissues of similar mitotic activity based on the EpiTOC2 model (Figure S2F). The physiological cause or consequence of this PMD hypomethylation in acinar cell biology warrants further investigation.

### MSA reveals dynamic 5-hydroxymethylation biology in human tissues

The standard array preparation uses bisulfite conversion, which does not discriminate 5-methylcytosine (5mC) from 5-hydroxymethylation (5hmC)^66^. To test if MSA is compatible with 5hmC profiling, hence producing a *ternary code* (5mC, 5hmC, and unmodified C) methylome, we employed a modified ACE seq protocol across the bulk human tissues^67^ (Methods). The derived 5hmC levels were globally anti-correlated with the proliferation rate of the tissue, being most abundant in neuron-enriched central nervous system tissues, followed by the kidney, heart, and liver, and lowest in the colon and lymph node (Figure S3A, S3B). Across chromatin states, 5hmC levels peaked in H3K36me3/H3K79me2 marked gene body enhancers and actively transcribed states (Figure 3A). Meta gene analysis showed a rapid depletion of 5hmC levels near the TSS, which rebounded and peaked in gene bodies (Figure S3C). To validate 5hmC measurements, we compared probe sets selected for tissue-specific 5hmC levels identified from publicly available 5hmC-Seal^39^ and hmC-CATCH^38^. While brain tissues had high 5hmC levels across most design groups, the non-brain tissues had the highest 5hmC in the designed tissue groups (Figure S3D).

**Figure 3:**
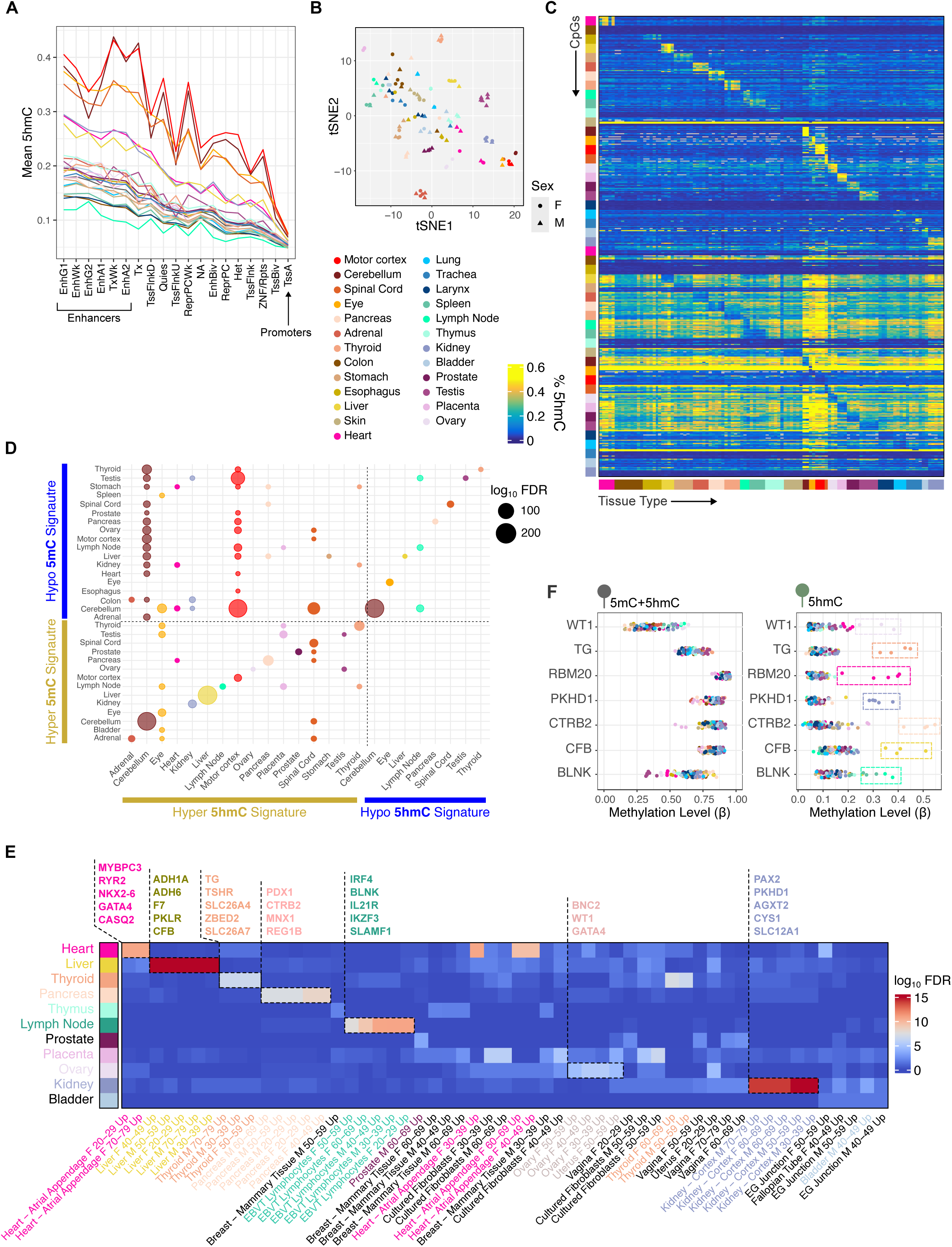
5hmC analysis of human tissues with MSA. (A) Line plot showing mean 5hmC levels across consensus ChromHMM states for each tissue type. (B) tSNE plot showing unsupervised clustering of bulk tissues profiled for 5hmC. (C) Heatmap showing representative one vs. all 5hmC signatures (rows) and the beta value across profiled tissues (columns) (D) Dot plot showing enrichment of 5modC tissue signatures in 5hmC tissue signatures. (E) Heatmap showing enrichment of genes linked to 5hmC CpG signatures for each tissue type in gene ontology sets from GTEx tissue-specific gene expression (columns) (F) Representative marker genes for seven tissue types showing 5modC (left) and 5hmC beta levels (right). Only 5hmC discriminates target tissue type.

Next, we expanded the tissue-specific 5hmC analysis to all probes. tSNE analysis of 5hmC profiles showed a separation according to tissue type (Figure 3B). Supervised analysis (Methods) identified dozens to thousands of tissue-specific 5hmC sites in most tissues (Figure 3C, S3E), the majority of which were associated with elevated 5hmC in the target tissue (Figure S3E). There were relatively few or no markers for skin (N=1) and colon (N=0), consistent with the low global 5hmC levels in these tissues.

Intriguingly, tissue-specific 5hmCs were highly enriched in the tissue-specific gain of 5mCs we identified via standard 5mod-C profiling in the matched tissue types (Figure 3D) and in an independent WGBS data set of sorted human cells^24^ (Figure S3F). Indeed, we found little overlap of tissue-specific 5hmC with tissue-specific loss of 5modC. As a result, the tissue-specific 5hmC was enriched in promoter states and, to a lesser extent, enhancers (Figure S3G), in contrast to the hypomethylation signatures we identified that were highly enriched in tissue-specific enhancers and transcription factor binding sites (Figure 2D, 2E).

Despite this lack of overlap, 5hmC still accumulated in a highly tissue-specific fashion. We tested the enrichment of genes in proximity to the 5hmC markers for each tissue type against GTEx tissue-specific RNA expression (Methods) and observed strong enrichment of the tissue marker genes identified by 5hmC in the corresponding tissue-specific RNA set from GTEx (Figure 3E). For example, liver-specific 5hmC linked to *CFB, PKLR,* and *ADH1A*, all genes that are specifically expressed in the liver^68^. Similarly, kidney 5hmC localized to *PKHD1, PAX2*, and *CYS1* which regulate kidney development and physiology^69,70^. At the probe markers for many of these tissue-specific genes, we did not observe hypomethylation colocalizing with 5hmC in the target tissue type; rather, we observed consistent hyper 5hmC across all the samples in the tissue group (Figure 3F). These results suggest that 5hmC may accumulate and remain stable in promoters and gene bodies to confer tissue-specific function, as opposed to existing as a transient byproduct of active demethylation pathways.

In contrast, hypomethylation occurs at more distal enhancers in the binding sites of tissue-specific transcription factors. These two modifications appear to be complementary in their localization and regulation of tissue-specific gene expression and cell identity. Future studies may elucidate the mechanisms by which 5hmC and 5mC are specifically targeted to their distinct locations in relation to tissue-specific genes.

### 5mC and 5hmC methylation biology in imprinting, aging, and sex specificities

To investigate methylation biology further, we analyzed constitutive methylation patterns not dictated by cell identity, across all the profiled tissues. Among the 128 tissues profiled, 13,633 probes were consistently unmethylated (β < 0.2), 5,012 were consistently methylated (β > 0.8), and 225 displayed intermediate methylation (β between 0.3–0.7; Figure 4A). Constitutively methylated CpGs were enriched in gene bodies, while unmethylated sites were predominantly found in CpG islands and transcription start sites (Figure S4A). Both categories were depleted in enhancer regions, which showed greater variability and are critical for tissue-specific regulation (Figure 2D).

**Figure 4:**
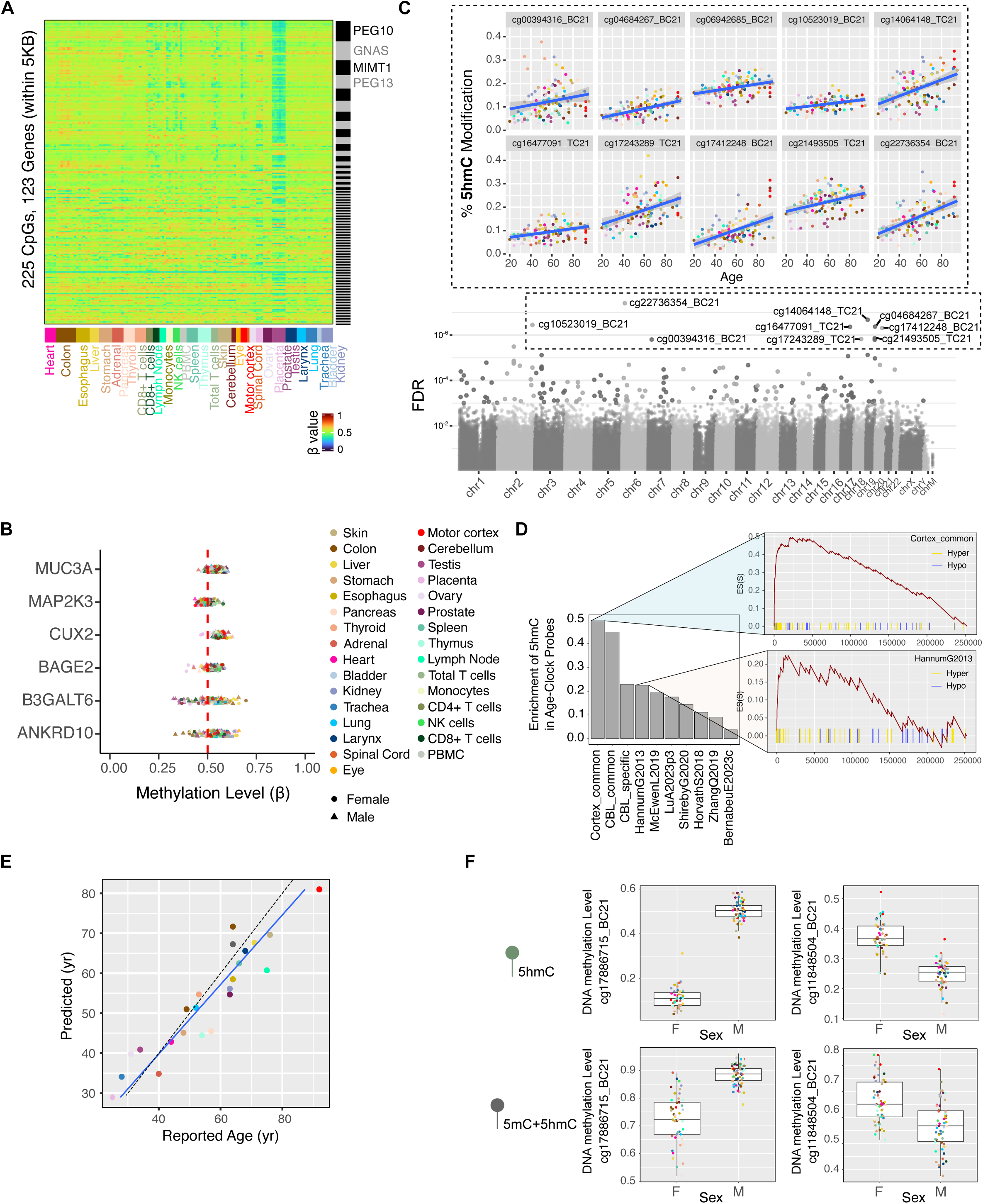
Epigenetic aging and mitotic history analysis of MSA profiled tissues. (A) Heatmap showing beta values over probes identified to be intermediately methylated (rows) across profiled samples (columns) (B) Mean beta values over intermediately methylated probes for six representative genes. The beta value patterns resemble those at known imprinting genes (C) Manhattan plot of the aging 5hmC EWAS (bottom) and scatter plots for representative age-associated 5hmC CpGs (top) (D) Set Enrichment scores for 10 epigenetic clocks. Epigenetic clock probes were tested for enrichment against a list of 5hmC probes, ranked according to the P value associated with aging. Representative panels on the right show clock probes enriching toward the top of the list of most significant 5hmC probes. (E) A scatter plot shows the relationship of reported age to predicted age on testing data for the multi-tissue clock trained to predict age using 5hmC. (F) Boxplots showing beta value distributions for 5modC and 5hmC over representative sex-specific autosomal CpGs

We linked intermediately methylated probes to genes within 5 kb, identifying 123 proximal genes. Notable genes with the most linked probes included known imprinting loci such as *PEG10*, *GNAS*, and *MIMT1*, which exhibit parent-of-origin expression regulated by methylation at imprinting control regions (ICRs) and differentially methylated regions (DMRs) (Figure S4B). Across 32 tissue types, the average methylations linked to these genes centered around 0.5 with minimal variability, except in the testis, which deviates due to sperm presence. Dozens of genes with proximal intermediately methylated probes displayed characteristics like ICR genes but are not to our knowledge currently documented as imprinted or monoallelically expressed (Figure 4B).

We further analyzed methylation and 5hmC patterns across aging using linear modeling (Methods) and identified thousands of age-associated CpGs, predominantly hypermethylated with age (Figure S4C). These CpGs were significantly enriched in PRC2 target regions, CpG islands, and bivalent chromatin (Figure S4D). Notably, 10 CpGs exhibited tissue-independent 5hmC increases during aging (Figure 4C). Set enrichment analysis revealed a strong overlap between age-associated 5mC CpGs and ranked 5hmC aging CpGs (Figure S4E), suggesting that some hypermethylation with age reflects 5hmC accumulation.

To further explore this, we assessed 20 epigenetic clocks with various degrees of age correlation in our 5mC datasets (Methods, Figure S4F). We found significant enrichment of clock probe sets in 5hmC aging probes, implying that clocks incorporate, to different degrees, 5hmC to estimate age (Figure 4D). Using 5hmC data alone, we trained a highly accurate model to predict chronological age (Figure 4E). These findings indicate that 5hmC is dynamic throughout the lifespan, and it alone can serve as a robust aging biomarker. Further investigation is needed to determine whether 5hmC accumulation results from spurious TET activity and if 5hmC age acceleration is associated with disease states.

Lastly, thousands of CpG sites showed sex-associated 5mC and 5hmC patterns, with 1,809 sites shared between the two modifications (Figure S4G, S4H). Most sex-associated CpGs were located on sex chromosomes, enriched in CpG islands and TSS chromatin states, likely reflecting sex-specific regulation of gene dosage (Figure S4I). Additionally, we identified 966 autosomal CpGs associated with sex for 5mC and 79 for 5hmC, some exhibiting differences as pronounced as those seen in X-linked CpGs (Figure 4F). The mechanisms underlying sex-specific methylation at autosomal loci and its potential role in regulating sex-specific expression and phenotypes remain to be explored.

### MSA methylomes reveal strong tissue contexts of human trait associations

Leveraging the trait association focus of MSA, we evaluated the capacity of MSA data to perform functional annotation of EWAS hits. In this analysis, we focused on the tissue context using the primary tissue profiles produced in this study. We first note that for the traits investigated in the curated studies, trait-associated probes are more often significantly enriched in enhancers and promoters^53^ but underrepresented in heterochromatic and repressive genomes (Figure S5A), consistent with their roles in transcriptional regulation. Traits characterized by genomic alterations (e.g., Down’s syndrome), cell proliferation (e.g., malignancy), and frequent toxin exposure (e.g., smoking) had distinct and recurring chromatin feature enrichment (Figure 5A). In contrast, complex disease traits, e.g., diabetes and Alzheimer’s disease, are varied in chromatin state enrichment across studies.

**Figure 5:**
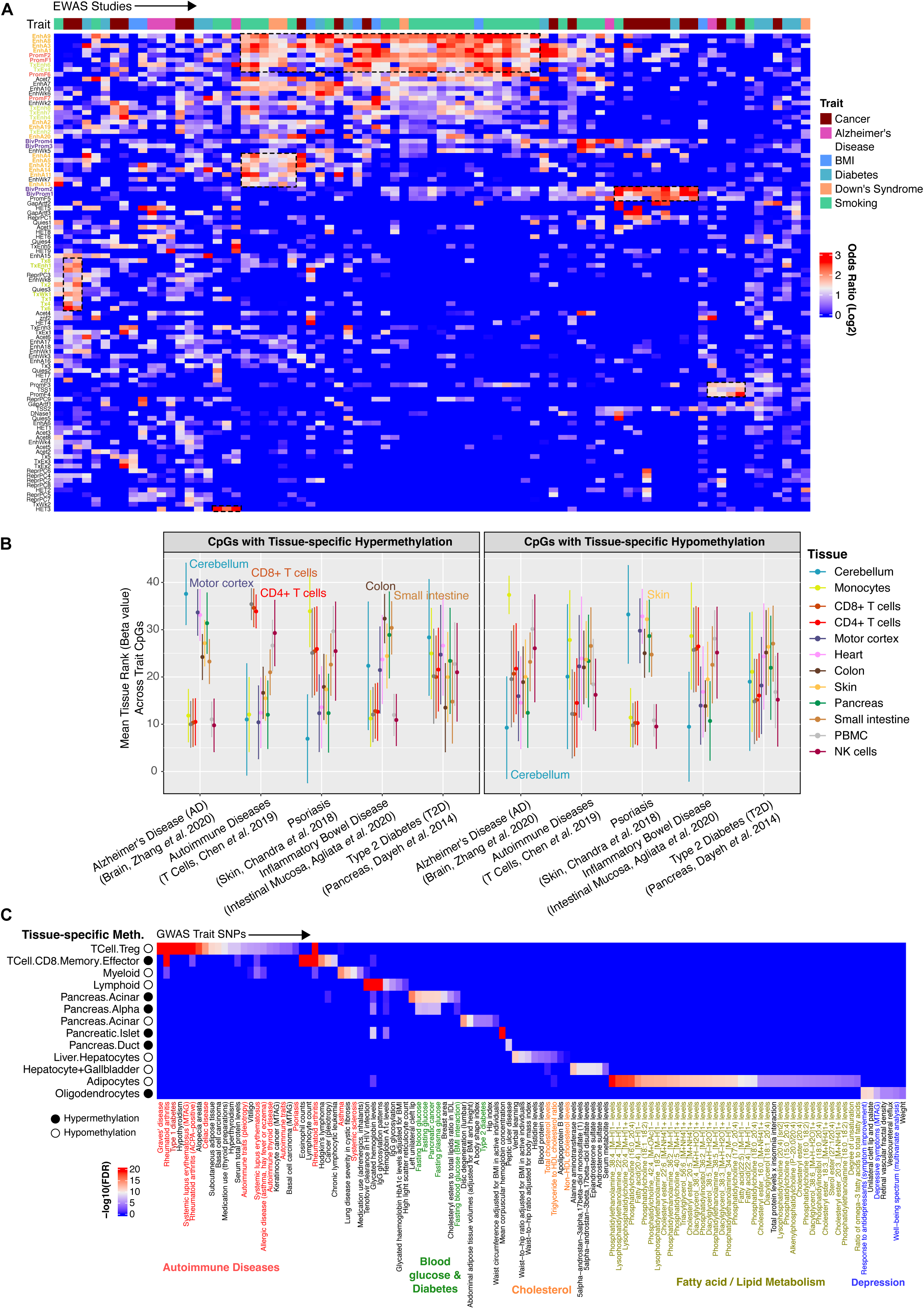
Tissue context of human trait associations. (A) Heatmap showing the enrichment of publicly available trait-associated probes across different genomic features (B) Distributions of mean beta value rank for each tissue type over trait-associated CpGs (C) Enrichment of a subset of designed tissue-specific methylation sets on MSA in the SNP sets associated with different GWAS traits

As expected, the enhancer and promoter-associated probes are more variably methylated across primary human tissue types (Figure S5B). To test whether such variation reveals the tissue context of each trait, we grouped CpGs by their associated traits and compared the methylation levels across tissue types (Figure 5B). An intriguing correspondence between the perceived tissue context and the methylation rank emerged. For example, CpGs associated with Alzheimer’s disease showed the most extreme methylation in brain tissues compared to other tissue types (Figure 5B). Sites with a putative positive disease effect size have the highest methylation readings in the brain, whereas sites with reduced methylation in diseases were least methylated in brain tissues. Similarly, probes associated with irritable bowel syndrome (IBS) were most methylated in the colon and small intestinal tissues. These results suggest a propensity of trait-associated CpGs to colocalize with differential methylations specific to the tissue that manifest the trait phenotype, underscoring the importance of tissue context when conducting EWASs.

We also investigated the extent to which GWAS variants colocalize with tissue-specific methylation. We tested the enrichment of trait-associated SNPs in the one-vs-all cell-specific methylation signatures on MSA (Methods). These analyses identified multiple genetic variants associated with a tissue-specific trait co-localizing with the methylation signature of the corresponding tissue type. For example, SNPs associated with blood glucose and diabetes were colocalized with methylation markers for pancreatic cell types, while cholesterol variants were localized to hepatocyte-specific methylations (Figure 5C). Diverse autoimmune disorders were enriched in CpG markers for regulatory T cells, which are involved in immune system homeostasis and autoimmune suppression^71^. Whether the genetic variants implicated in these diseases directly impact nearby tissue-specific methylation to perturb gene expression and function requires follow-up studies.

### MSA detects inter-individual methylation variation at EWAS trait sites

To date, thousands of traits have been analyzed in EWAS studies using peripheral whole blood, a clinically accessible tissue source that provides sufficient DNA for array-based analysis. To explore immune cell dynamics and evaluate the array’s capacity for detecting interindividual variation, we analyzed 64 whole blood samples from anonymous donors using MSA. The MSA design included some major epigenetic clocks (Figure S6A), and we verified that we could accurately predict age using the multi-tissue Horvath clock^72^ on the tissues we previously profiled (Figure S6B). The Horvath clock and a sex prediction model (Methods) applied to the whole blood samples revealed a broad age range (8.7–58.4 years) and a sex distribution of 14 females and 50 males (Methods, Figure S6C).

Cell composition explains most bulk-tissue epigenetic variations. To analyze interindividual cell composition variation using DNA methylation, we first benchmarked computational deconvolution on MSA-based methylation profiles of sorted immune cells. As expected, predicted sorted immune cells contained >90% of the matching cell type, consistent with standard purification yields (Methods, Figure 6A). Then, we applied the same deconvolution strategy to whole-blood DNA methylomes. The results yielded estimates aligned with prior literature (Figure 6B; mean estimates: Neutrophils 61%, CD4T 14%, CD8T 9%, Monocytes 7%, B Cells 6%, NK 3%). Principal component analysis showed immune cell proportions, along with sex, explained the greatest variance in the data set (Figure 6C, S6B). To examine immune cell composition dynamics, we regressed cell type proportions on predicted age and sex. We found that aging was associated with a significant decrease in CD4+ T cells and an increase in neutrophils (Figure 6D). Sex differences revealed higher CD8+ T cell proportions and lower NK cell proportions in females (Figure 6E).

**Figure 6:**
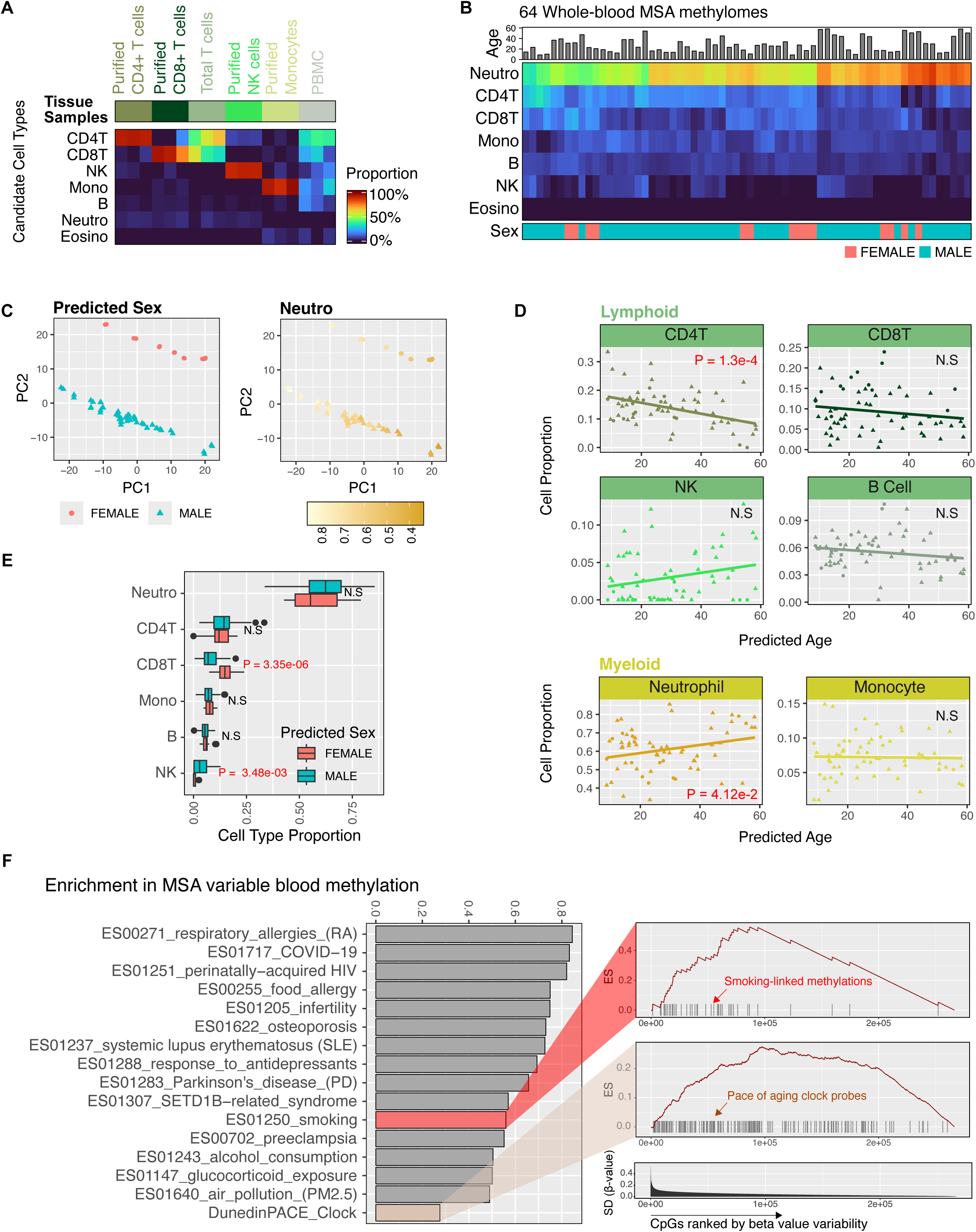
Immune cell composition and interindividual whole-blood methylation variation. (A) Validation of immune cell deconvolution using sorted immune cell methylation profiles. (B) Immune cell proportion estimates in 64 whole blood methylomes. (C) Principal component analysis shows that immune cell proportions and sex explain the largest variance across the dataset. (D) Age-associated immune cell composition dynamics: CD4+ T cell proportions significantly decrease with age, while neutrophil proportions increase. (E) Sex differences in immune cell composition. (F) Enrichment of previously reported EWAS traits in CpG sites with high inter-individual methylation variation.

To further assess interindividual variations, we ranked autosomal probes by standard deviation across individuals. Using a set enrichment framework (Methods), we observed that sites with inter-individual methylation variation are significantly enriched in EWAS traits previously reported by blood-based EWASs, including immune system disorders and other environment-related traits (e.g., smoking and alcohol consumption) (Figure 6F). The new MSA probe designs showed a similar distribution of inter-individual variations compared to legacy probes, suggesting an expanded capacity for detecting blood-based methylation-trait links (Figure S6C). While we could not directly correlate methylation with phenotypic traits in our data set, the results demonstrate that MSA detects methylation variations associated with various physiological outcomes identified in prior studies.

## DISCUSSION

The Infinium DNA methylation BeadChip is a broadly used and accessible assay in human population studies. It has enabled trait association discoveries and predictive models such as epigenetic clocks, risk scores, and disease classifiers. Previous Infinium BeadChips have been designed to target genomic features, such as gene promoters, gene bodies, and cis-regulatory elements. While methylation variation at these genomic features is indeed associated with human traits, evenly covering genomic elements is not as economical for trait screening applications as in discovery and hypothesis generation settings.

The existing methylation-based screening of most human traits requires relatively few loci. For instance, the Horvath clock for chronological age used 353 CpGs^72^. Other epigenetic clocks use feature numbers ranging from a few CpGs to ten thousand CpGs^73^, which are much smaller in number than existing Infinium array capacities^43^. The feasibility of such minimalistic approaches has also been established in cancer classification^48^ and cell type deconvolutions^74^ and demonstrates high inference precision. The development of MSA can be seen as a balanced approach to DNA methylome-based trait screening, prioritizing only the probe sets that link to diverse traits and high-confidence prediction models for the benefit of profiling larger human populations.

While legacy probes were incorporated for their established trait associations, the enhanced scalability of MSA may facilitate the repositioning of these probes for novel associations. Historically, populations of European descent have been overrepresented in EWAS studies, potentially overlooking disease-relevant associations in more diverse demographics. Re-examining these associations in larger and more balanced cohorts will be imperative to dissecting the complex interplay of genetic and environmental influences on disease phenotypes. The legacy probe designs chosen for inclusion in MSA are also frequently associated with multiple traits, implying that multiple physiological or environmental stimuli can converge on similar epigenetic programs. Future studies may elucidate whether these shared signatures represent common inflammatory or homeostatic pathways that are similarly disrupted and whether additional, currently under-studied disease states converge on the same loci.

Besides offering a balanced approach in trait screening, MSA also represents an upgrade of Infinium array content to bridge deep high cell-type resolution profiling and cost-effective population screening. While offering greater cell type variation and genome-wide details, single-cell methylome profiling cannot be scaled to population settings. MSA is designed to translate the cell type-specific knowledge from single-cell and bulk whole-genome methylome profiles for use in the population setting.

Computational cell-type deconvolutions are powerful methods for interrogating tissue composition variation in development and disease. The expanded cell-specific CpG markers and refined annotation in MSA enhance deconvolution granularity compared to EWAS studies based on previous Infinium platforms. For example, the commonly used CETS algorithm for estimating brain cell proportions estimates NeuN+:NeuN-proportions without predicting trait-relevant subtypes^75^. We designed cell-specific probes discriminating 174 unique cell types (82 brain cell types, 51 pan tissue, 41 blood) and anticipate that these markers will enable high-resolution deconvolution, augmenting the study of selectively vulnerable or rare cell populations in complex diseases and tissue types. Our results and other recent work have also identified an enrichment of genetic variants associated with complex traits within cell-specific DMRs^22^. It is not clear the extent to which methylation changes in these cell-specific DMRs may perturb the functioning of the disease-relevant cell types. We anticipate that MSA will permit such investigations.

Previous efforts have established the compatibility of Infinium arrays with other base conversion protocols, such as Tet-assisted bisulfite conversion, to profile 5hmC modifications^76,77^. Our analysis suggested that the new MSA array is compatible with the tandem bisulfite-A3A conversion for 5hmC profiling. We applied the 5hmC profiling to neuronal and peripheral human tissues. The tissue-specificity mirrors previous sequencing-based 5hmC profiles, suggesting the feasibility of using methylation arrays to implement 5hmC profiling in large sample sets. Our data also underscores the high cell type specificity of 5hmC signals, which are often distinct but complementary to cell-specific hypo 5modC and could be additionally used to trace cell identity and tissue composition changes. Over aging and across tissues, we identified dynamic 5hmC variations that are strongly linked to tissue specific gene expression and aging prediction models.

As a first application, our analysis was limited in validating the trait-associated probes selected due to limited metadata availability. However, we found that probes associated with some traits in the literature were variably methylated in the corresponding tissue types we profiled or had a strong tissue context according to the beta value rank by tissue type (Figure 5). Attempting to design a consolidated array, we were also limited in the number of the CpG sites we could include and thus genomic feature and trait coverage. As more WGBS and array-based methylomes are generated, future designs may refine the most relevant trait and cell type-implicated CpG sites to maximize screening and discovery power most economically.

## CONCLUSION

We systematically developed, benchmarked, and applied MSA, a novel Infinium BeadChip assay consolidating trait-associated probes from the extensive EWAS literature, single-cell and bulk whole genome methylome profiles. Our benchmark revealed MSA as an accurate, reproducible, scalable, next-generation Infinium human methylation BeadChip targeting trait discovery in population settings. Our first application uncovered the cell type context of human EWAS and GWAS discoveries and dynamic 5hmC association in peripheral tissues. We anticipate MSA to be a valuable tool for methylation screening in large human populations for trait associations and broadly dissecting the cell-type-specific mechanisms of human diseases.

## METHODS

### CpG Probe Selection

#### Probe designability

We aligned unmethylated and methylated probe sequences to the hg38 genome using the BISCUIT tool suite^78^. To identify uniquely mapping sequences, subsequences of 30,35,40 and the entire 50nt probe sequence were aligned, and only probe designs where all subsequences had mapping quality >20 for both the methylated and unmethylated allele were considered. For these 19,253,974 uniquely mapping CpGs, design scores reflecting hybridization efficiency and melting temperature were computed, and 13,891,035 CpGs with design scores > .3 were retained. Any probe sequence that contained common SNPs (dbSNP Build 151)^79^ within 5nt of the 3’ end was removed. Sequences with more than six additional CpGs were also removed to prevent hybridization interference due to variable methylation of neighboring CpGs. 9,993,793 CpGs remained from this preprocessing (“Designable Probes”), from which all array content was subsequently selected. When possible, high-quality probes (design score >= .6) were prioritized.

#### Cis-regulatory elements

Human GRCh38 candidate cis-regulatory element (CRE) annotations were downloaded from the ENCODE Project Consortium^80^ and intersected with designable CpG sites. The methylation range for each CpG was computed across sorted immune^52^ and pan tissue^24^ cell types. CpGs that did not show a range > .4 were filtered out. The remaining CpGs were grouped by CRE type and sorted by methylation range. 30,000 CpGs total were sampled with a bias toward enhancer elements (dELS: 64%; pELS: 21%; CTCF Only, CTCF-bound:11%; PLS:2%; DNAse-H3K4me3:2%).

#### Monoallelic/intermediate methylation

180 bulk adult normal WGBS samples (Table S2) were analyzed to identify candidate monoallelically methylated CpG sites. Autosomal CpGs with minimum coverage of 20 reads and mean methylation >.3 and <.7 across 140 of the 180 samples were considered intermediate methylation and intersected with the designable probe list. 207 pan-tissue sorted cell WGBS methylomes from Loyfer et al ^24^ were also analyzed for intermediate methylation, and designable CpGs with mean methylation >.3 and <.7 across 180 of the 207 samples were selected.

#### XCI-linked CpGs

76 high coverage (>20 million CpGs) normal female WGBS samples (Table S2) were analyzed to identify X-chromosome CpG sites with intermediate methylation across samples (0.3 < methylation < .7). An additional 95 normal male WGBS samples were analyzed to identify X chromosome CpG sites fully unmethylated (< .3 methylation across 50 samples) or fully methylated (>.7). The CpG sites intermediately methylated in female samples but unmethylated or fully methylated in male samples were intersected with the high-quality probe list.

#### Cell type-specific methylation

BED/bigWig files for single cell brain^49–51^, sorted pan tissue^24^, and sorted immune cell WGBS data^52^ were downloaded and used for marker identification. To reduce the sparsity of single-cell brain data, pseudo bulk methylomes were generated by averaging methylation over the cell type labels obtained by unsupervised clustering analysis previously reported. One vs. all comparisons were performed across major cell type groups and hierarchically within major groups to identify subtype markers. Wilcoxon rank sum testing was performed between the target and out groups at each CpG site to identify cell-specific markers. Designable CpG sites with an AUC = 1 and a delta beta >= .3 between the in and out groups were selected, and markers were capped at 80 CpGs per cell type contrast. Hyper and hypomethylated signatures were balanced when possible.

#### 5hmC

5hmC-Seal^39^ and hmC-CATCH^38^ 5hmC peaks were downloaded. Genomic intervals were intersected with the designable CpG list. For 5hmC-Seal data, the 5hmC CpG signal was treated as a binary value (1 if within a significant peak, 0 if not). For hmc-CATCH data, the peak coverage was applied to CpGs within the peak, and samples were scaled according to the total coverage. Tissue-specific 5hmC sites were identified as previously described for the WGBS data. To identify 5hmC sites along a continuum of tissue specificity, the top 10K most highly covered CpGs in each sample from the hmC-CATCH data^38^ were collected and binned according to the frequencies the CpG was in the top 10K across the 60 samples. 11 bins of 5 tissue count intervals (e.g., 1-5, 6-10,…, 55-60 tissues) were sampled equally, with sampling capped at 200 CpGs per bin.

#### Cell-specific CpH methylation

Genes with cell-specific mCH methylation were downloaded^49^, and the top ten genes with the highest AUROC were selected for each cell type. Gene coordinates were intersected with CAC cytosines, the most prevalent mCH context found in neurons. 20 cytosines were sampled from each gene for each cell type.

#### DNA methylation-gene expression correlations

Matched WGBS / Gene expression data from the Roadmap Epigenomics Mapping Consortium were used to compute the Spearman correlation between CpGs in the high-quality designability list and genes within 10KB of the CpG. CpGs were then ranked by the *P*-value of the correlation, standard deviation and expression levels of the gene, and absolute value of the correlation. The top 2,500 CpGs negatively correlated with the expression of the linked gene, and the top 2,500 positively correlated CpGs were selected. TCGA normal tissues^81^ were also analyzed to identify correlated linked CpG-Gene pairs. CpGs with a correlation >= 0.6 or <= −0.7 and a *P*-value < .05 were additionally included (901 positively correlated, 1,620 negatively correlated).

#### DNA methylation-chromatin accessibility correlations

Matched DNA-chromatin accessibility data were downloaded from Luo *et al.* 2022^49^, and Spearman correlations were computed between the accessibility peaks and CpG methylation sites. Correlations with *P*-values < .05 and |Spearman’s rho| > .5 were selected, and the CpGs intersected with the high-quality designability list.

#### CoRSIVs

Genomic coordinates for CoRSIVs were downloaded^82,83^ and intersected with high-quality designable probes.

#### Solo-WCGW in partially methylated domains

CpGs in the WCGW context (flanked by A or T) in common PMDs were downloaded from Zhou *et al.* 2018^23^ and intersected with high-quality designable probes. This subset was further intersected with CpG islands, and 6,000 probes were randomly sampled.

#### meQTLs

meQTL data was downloaded from the GoDMC database^30^, and CpGs were ranked according to the number of times a CpG was associated with a meQTL. The top 10K CpGs were selected. An additional 20K meQTLs were randomly sampled from Hawe *et al.* 2021^29^.

#### Imprinting-associated DMRs

Differentially methylated regions associated with monoallelically expressed genes were downloaded from Skaar *et al.* 2012^84^ and lifted to GRCh38 coordinates. The DMRs were intersected with the designable probes list.

#### Y-linked genes

180 high coverage (>20 million CpGs) Human WGBS samples (Table S2) were analyzed to identify variably methylated Y-linked genes. The Y chromosome CpGs were intersected with designable probes and subsequently intersected with all Y chromosome genes (GENCODE V39). The variance across the 180 samples was computed at every remaining CpG site. For each gene, the top 20 most variable probes were selected.

#### Human trait associations

1,067 EWAS studies were curated from the literature and EWAS databases (EWAS catalog^33^, EWAS atlas^34^). A subset of high-priority studies was identified according to sample number and statistical significance, diversity of trait coverage, citation number, and the journal impact factor. All probes, or the top 2500 most significant probes, were selected from high-priority studies. The top 100 most significant probes were selected from all remaining curated studies. Study titles and trait annotations were queried for regular expressions to consolidate all selected studies/traits into 16 major trait groups.

### Sample Preparation

#### Tissue dissection

Fresh frozen tissue samples were obtained from the Cooperative Human Tissue Network (CHTN), and 30-50mg of tissue were dissected on dry ice.

#### Cell line culture

GM12878, K562 (CCL-243), LNCaP (CRL-1740), and HCT116 (CCL-247) cells (Source 1) were obtained from American Type Culture Collection (ATCC, Manassas, VA, USA). 1-4 x 10^6 cells were plated and cultured for 6 days with fresh media added 2-3 days. K562 cells were cultured in Iscove’s Modified Dulbecco’s Medium (30-2005, ATCC), 10% Fetal Bovine Serum (FBS) (45000-736, Gibco), and 1% penicillin/streptomycin (15140122, Gibco). LNCaP cells were cultured in Roswell Park Memorial Institute Medium (RPMI-1640) (30-2001, ATCC), 10% FBS, and 1% penicillin/streptomycin (15140122, Gibco). GM12878 cells were cultured with RPI-1640 (72400047, Invitrogen), and 15% Fetal Bovine Serum (Gibco, 45000-736), 1% GlutaMAX™ (Gibco, 35050061), and 1% penicillin/streptomycin (15140122, Gibco). HCT116 cells were cultured in McCoy’s 5a medium modified (ATCC,30-2007), 10% Fetal Bovine Serum (FBS) (45000-736, Gibco), and 1% penicillin/streptomycin (15140122, Gibco). All cells were maintained in a 37°C incubator with 5% CO2 and cultured at a 75 cm2 culture flask (Fisher, BD353136)

#### DNA extraction

Genomic DNA was extracted from 30-70 mg of tissue or 5.0 x 10^6 cells for Source 1 cell lines using commercially available QIAGEN QIAamp Mini Kit (QIAGEN, 51304), following the manufacturer’s protocol. DNA was quantified using a Qubit 4 Fluorometer (Invitrogen). For Source 2 and Source 3 cell lines, genomic DNA was purchased from BioChain Institute (HeLa - #D1255811, Raji - #D1255840, Jurkat - #D1255815, MCF7 - #D1255830, K562 - #D1255820)

#### Immune cell purification

Sorted immune cells were purified by the Human Immunology Core at the University of Pennsylvania following STEMCELL Technologies RosetteSep Enrichment Cocktail protocols (https://cdn.stemcell.com/media/files/pis/10000000545-PIS_02.pdf). PBMCs were isolated using a Lymphoprep ficol layer.

#### Methylation titration controls

10 ng of fully methylated human blood (Thermo Scientific, SD1131) and Jurkat (Thermo Scientific, SD1121) genomic DNA were amplified using the Repli-g Mini Kit (QIAGEN, 150023) according to the manufacturer’s protocol. Following quantification with a Qubit 4 Fluorometer, 500ng of unamplified and amplified DNA were combined for the 50% control. Human pre-mixed calibration standards (0,5,10,25,50,75,100%) were purchased from EpigenDx (EpigenDx 80-8060H_PreMix), and 200ng / titration was used for testing.

#### EM sequencing of cell line DNA

Genomic DNA from the GM12878, K562, and HCT116 cell lines were extracted according to the QIAGEN QIAmp Mini Kit Protocol. The three samples were then mechanically sheared to 300 base pairs using the M220 Focused-ultrasonicator (Covaris, 500295) and methylated lambda control DNA. 200ng of each sample was enzymatically converted using the NEBNext® Enzymatic Methyl-seq Kit (NEB, E7120) with the manufacturer’s protocol. The samples were then indexed during PCR amplification during PCR amplification using EM-Seq™ index primers (NEB 7140). The indexed libraries (200 ng each) were pooled and used as input for the Twist NGS Methylation Detection System for target enrichment. A pre-hybridization solution of blockers and enhancers was created to prepare the pool for hybridization (Twist Bioscience, 104180). The DNA was hybridized with the Twist Human Methylome Panel (Twist Bioscience, 105520), and the targets were bound with streptavidin beads (Twist Bioscience, 100983), followed by a post-capture amplification. The enriched libraries were sequenced to 20X on the Illumina Novaseq 6000 PE150 platform.

#### 5hmC profiling

Using the EZ DNA Methylation Kit (Zymo Research, D5001), 500 ng of each sample was bisulfite converted and purified following the manufacturer’s protocol. The samples were then denatured with DMSO at 95°C for 5 minutes and snap-cooled on dry ice. The samples were deaminated using APOBEC3A (A3A) purified following previously published protocol^85^ over 2 hours at 37°C. After incubation, the samples were purified using the Oligo Clean and Concentrator Kit (Zymo Research, D4060), following the manufacturer’s protocol. Two cycles of whole genome amplification were performed using 50 U of Klenow Fragment (3’→5’ exo-) (NEB, M0212M), dNTP solution mix (Bio-Rad, #1708874), and Random Primer 6 (NEB, S1230S). The samples were finally purified using AMPure XP Beads (Beckman Coulter Life Sciences, A63881).

### MSA Data Analysis

#### Data preprocessing

All data preprocessing was done using the *SeSAMe* R package (version 1.22.0)^56^. A manifest address file was generated using the MSA manifest available at https://github.com/zhou-lab/InfiniumAnnotationV1/raw/main/Anno/MSA/MSA.hg38.manifest.tsv.gz and the *sesameAnno_buildAddressFile* function. Beta values were extracted from raw IDAT files using the *openSesame* function with the built address file and default parameters. Probe detection rates were obtained using the *probeSuccessRate* argument with the *openSesame* function. One sample with probe detection rates < 0.7 was excluded from analyses.

#### Trait enrichment testing

2,398,372 EWAS hits were curated from the literature and EWAS databases^33,34^ and used as a background for enrichment testing. Traits were annotated to 16 major trait groups by searching for regular expression terms relevant to the trait group within the study or trait descriptions. The odds ratio enrichment in these trait groups was computed for 3 query sets: 1) EPICv2 probes, retained MSA probes from prior Infinium platforms, and a random set of probes equal in size to the retained MSA probes. log2 odds ratio was plotted for each platform across trait groups. For testing the enrichment of MSA and EPICv2 probes in total trait associated probes, all EWAS probes were rank-ordered according to how many traits the probes associated with. The MSA and EPICv2 probes were each tested as a query against the ranked probe list using a modified gene set enrichment approach^86^ using the *knowYourCG* R package (version 1.0.0).

#### Gene linkage and ontology analysis

The MSA and EPICv2 manifests were downloaded (https://github.com/zhou-lab/InfiniumAnnotationV1/raw/main/Anno/), and probe coordinates expanded 1500bp upstream of the probe start site. The manifests were then intersected with GENCODE.v41 GTF files to identify linked genes. Gene ontology testing was performed for protein-coding genes using Enrichr^87^. The GO Biological Process gene set was queried. For CpH probe-linked genes, only genes with a minimum of 2 probes per gene were analyzed.

#### Sample reproducibility and accuracy

Pearson correlation coefficients were computed across cell line samples. Correlation matrices were plotted in heatmaps. For pairwise replicate comparisons, beta values were first binarized as 1 if beta > 0.5 and 0 if beta < 0.5. F1 scores for the binarized vectors were computed using the MLmetrics package (1.1.3).

#### Cell deconvolution

Reference-based cellular deconvolution for sorted immune cells and whole blood samples was performed using the EpiDISH R package^88^ (version 2.18.0) with the robust partial correlations (RPC) method. The centDHSbloodDMC.m matrix provided within the package was used as a reference for sorted immune cell deconvolution. For bulk tissue cell type inference, a reference for one vs. all cell-specific CpGs was created from Loyfer et al. 2023^24^ as previously described and deposited to the CytoMethIC github repository (https://github.com/zhou-lab/CytoMethIC_models/). Cell proportion scores were computed with the *cmi_predict* function from the CytoMethIC package (Version 1.1.1)

#### Identification of tissue-specific markers

One-vs-all tissue type comparisons were performed for sorted immune cells and bulk tissues. Wilcoxon rank sum testing between the target and out-group was performed at each CpG site. CpGs with NA values in >10% of the target group or >50% of the out group were excluded. The AUC for discriminating between the target and the out-groups was computed. Only CpGs with a delta beta >20% and AUC >= .8 were selected as cell markers. For visualization, the top 50 hypo and hypermethylated CpGs sorted by AUC and delta beta were selected for each tissue type. For 5hmC samples, the same analysis was performed, and a delta beta of 5% was used as a threshold for marker identification.

#### Tissue-specific CpG - transcription factor binding site analysis

BED files containing TFBS peaks were downloaded from ReMap 2022 (https://remap.univ-amu.fr^89^). The peaks for each transcription factor were intersected with all MSA CpGs to create CpG-TFBS links. Tissue signatures were tested for enrichment in the TFBS CpG sets using Fisher’s exact test with all MSA probes as the background.

#### Enrichment testing in chromatin states

Enrichment testing in chromatin states for all probe sets in manuscript was performed using the *knowYourCG* R package (version 1.0.0) with the chromHMM knowledgebase set and *testEnrichment* function.

#### Tissue-specific CpG marker validation enrichment testing

BED/bigWig files for single cell brain^49–51^, sorted pan tissue^24^, and sorted immune cell WGBS data^52^ were downloaded and used for marker identification. To reduce the sparsity of single-cell brain data, pseudo bulk methylomes were generated by averaging methylation over the cell type labels obtained by unsupervised clustering analysis previously reported. One vs. all comparisons were performed across major cell type groups and hierarchically within major groups to identify subtype markers. Wilcoxon rank sum testing was performed between the target and out groups at each CpG site to identify cell-specific markers. CpG sites with an AUC > .95 and a difference in beta value > .5 between the in and out groups were selected to generate marker lists for each cell type and intersected with MSA probes. The 5modC and 5hmC tissue signatures identified from MSA profiled tissues were tested for enrichment in the marker lists using Fisher’s exact test with all MSA probes as the background.

#### Nearest neighbor analysis

Nearest neighbor analysis was performed using deep WGBS data^24^ to identify neighbor genomic coordinates on MSA for non-retained EPIC probes. The WGBS data was subset for the MSA probe genomic coordinates and reference graphs were constructed using the *nnd_knn (k=50 neighbors)* function from the *rnn_descent* R package (version 0.1.6). The graph was then queried using the EPIC probe genomic coordinates from the WGBS data using the *graph_knn_query* function. For each CpG, the neighbor in the reference graph with the lowest Euclidean distance was recorded. We additionally computed the Euclidean distance between every EPIC probe and the nearest genomic neighbor on MSA. The final CpG with the lowest Euclidean distance was retained. To test the performance of neighbor probes in classifying tissue type, we used an EPIC tissue prediction model from the *CytoMethIC* R package (version 1.1.1) and removed all probes from the model that were retained on MSA. For remaining EPIC-only probes, we substituted the neighbor beta values from the MSA methylomes to compute the tissue inference.

#### Tissue-specific CpG – marker gene enrichment testing

CpG signatures for each tissue type were linked to genes +/- 10KB from the CpG site (GENCODEv19). The resulting gene sets for each tissue type were tested for enrichment against the HumanGeneAtlas^90^ downloaded from Enrichr^91,92^ and the top 5 most enriched ontology terms (FDR < .05) for each tissue type’s gene sets were selected for network graphing in Cytoscape version 3.9.1 using the log2 odds ratio for edge weights and an edge weighted spring embedded layout.

#### Epigenetic clock estimation

730 TCGA normal tissues profiled on the HM450 array were used to assess the impact of missing probes on epigenetic clock estimation. The full clock probes, and the subset represented on MSA were both tested, and the predictions compared (Fig S1O). For MSA profiled tissues, the probe suffixes were removed and duplicate probes averaged. All age estimates were computed with the *DNAmAge* function from the *methylclock* package (version 1.8.0)^93^ using default parameters. HypoClock and EpiTOC2 mitotic rate estimates were computed by tissue type group using the data and code provided by the authors at https://zenodo.org/records/2632938. Placental tissues were excluded.

#### Sex prediction

Sex for anonymous whole blood donors was inferred using the *cmi_predict* function from the *CytoMethIC* R package (version 1.1.1) using the sex associated CpGs from the models represented on the MSA array. This model generates a sex score by averaging the difference between male associated hyper and hypo methylation over known sex associated CpGs.

#### Linear modelling

Linear modelling for age and sex associated 5modC and 5hmC was performed using the *DML* function from the SeSAMe package^56^ version 1.22.0, covarying for tissue type (CpG ∼ Age + Sex + Tissue). P-values were adjusted for multiple comparisons using the FDR method and CpGs with FDR < .05 for age and sex were considered for further analysis. Testis and placenta excluded. For analysis of whole blood methylomes, cell type proportions from deconvolution analysis were regressed on epigenetic age and sex using linear models.

#### Set Enrichment Analyses

All set enrichment analyses were performed using the *testEnrichmentSEA* function from the *knowYourCG* package R package (version 1.0.0). For testing epigenetic clock probes against 5hmC age probes, epigenetic clock probes were downloaded from the *dnaMethyAge* R package (https://github.com/yiluyucheng/dnaMethyAge) and tested against the ranked list of age associated 5hmC probes, sorted according to P value from the 5hmC ∼ Age + Sex + Tissue EWAS. The top 10 most enriched clocks were plotted. For variable blood methylome analysis, autosomal probes were ranked according to the standard deviation across the 64 whole blood samples. EWAS trait CpGs^33,34^ were tested as queries against the variable probe list.

#### 5hmC age clock

For each tissue type, 80% of samples were randomly selected for training and the remaining 20% used testing. For feature selection, the top 1000 CpGs according to P value were selected from the 5hmC aging EWAS. An elastic net regression model was trained to predict age from the 5hmC beta values using the cv.glment function (alpha=0.5, nfolds=10) from the *glmnet* package (Version 4.1-8).

#### Analysis of EWAS hit chromatin state contexts

Each set of EWAS trait probes in the curated studies was tested for enrichment in 100 full-stack ChromHMM chromatin states ^53^ using Fisher’s exact test. The total pool of curated EWAS hits was used as a background set. The number of traits-chromatin state associations with FDR < .05 was computed for each chromatin state and plotted. 17 studies representing 6 trait groups were selected, and the enrichment across chromatin states was plotted in heat maps.

#### Chromatin context analysis of EWAS methylations

The standard deviation of all probes was computed using the tissue methylomes generated on MSA and sorted to create a ranked probe list. Selected full-stack ChromHMM states were intersected with the list of total EWAS hits and tested as queries against the ranked probe list using a modified gene set enrichment approach^86^ using the *knowYourCG* R package (version 1.0.0).

#### Tissue context analysis of EWAS methylations

For each set of EWAS trait probes in the curated studies, we computed the standard deviation of the probes using the tissue methylomes we generated using MSA. Trait sets were sorted according to the average standard deviations, and the most variable traits were selected for further analysis. In these traits, the rank for each sample was computed according to beta values. The mean rank of each tissue type group was computed for every CpG in the trait, and the distributions of ranks for each tissue type were plotted.

#### GWAS co-localization with tissue-specific methylations

GWAS summary statistics were downloaded from the NHGRI-EBI GWAS catalog^94^ (version 1.0.2.1). The top 3000 unique disease/trait categories with the most SNPs were grouped and tested as independent queries against each one-vs-all tissue/cell-specific CpG set from the curated lists incorporated into the final MSA design. SNPs and CpG sites were expanded by 5kbps in upstream and downstream directions, and genomic interval overlaps were computed using the *IRanges* package (version 2.36.0). The total number of CpG intervals for all tissue signatures was used as a background set, and Fisher’s Exact test was performed for enrichment testing.

## ACKNOWLEDGEMENTS

The authors thank Lynn Chen, Max Eldabbas, and Emileigh Maddox of the Human Immunology Core at the Perelman School of Medicine at the University of Pennsylvania for assistance with immune cell purification. The HIC is supported in part by NIH P30 AI045008 and P30 CA016520. HIC RRID: SCR 022380. The authors thank the NCI Cooperative Human Tissue Network (CHTN) for providing human tissue samples. Other investigators may have received specimens from the same tissue specimens. RRID: SCR_004446.

## AUTHOR CONTRIBUTIONS

Conceptualization, W.Z., N.R., R.P., and D.C.G.; Methodology, D.C.G., W.Z.; Formal Analysis, D.C.G., and W.Z.; Investigation, D.C.G., C.C, E.K., S.L., M.H., M.K. S.G., A.P., B.B., M.W, and L.M.; Resources, R.K., J.B.P., M.H., M.K., Q.Z., S.G. and A.P.; Writing – Original Draft, D.C.G. and W.Z.; Writing – Review & Editing, D.C.G., W.Z., NR, RP and RMK, Funding Acquisition, N.R., R.P. and W.Z.; Supervision, W.Z.

## Availability

Informatics for MSA data preprocessing and functional analysis is available in the R/Bioconductor package *SeSAMe* (version 3.22+): https://bioconductor.org/packages/release/bioc/html/sesame.html

The complete MSA manifest, design criteria, technical, human trait, and functional annotations are available at https://zwdzwd.github.io/InfiniumAnnotation

The generated human cell line, primary tissue 5mC and 5hmC methylome profiles (N=676), and EM-seq data are available in the Gene Expression Omnibus with accession GSE264438 and GSE267407.

## REFERENCES

1. Bird, A. (2002). DNA methylation patterns and epigenetic memory. Genes Dev. 16, 6–21. 10.1101/gad.947102.

2. Chen, R.Z., Pettersson, U., Beard, C., Jackson-Grusby, L., and Jaenisch, R. (1998). DNA hypomethylation leads to elevated mutation rates. Nature 395, 89–93. 10.1038/25779.

3. Greenberg, M.V.C., and Bourc’his, D. (2019). The diverse roles of DNA methylation in mammalian development and disease. Nat. Rev. Mol. Cell Biol. 20, 590–607. 10.1038/s41580-019-0159-6.

4. Flanagan, J.M. (2015). Epigenome-wide association studies (EWAS): past, present, and future. Methods Mol. Biol. 1238, 51–63. 10.1007/978-1-4939-1804-1_3.

5. Locke, W.J., Guanzon, D., Ma, C., Liew, Y.J., Duesing, K.R., Fung, K.Y.C., and Ross, J.P. (2019). DNA methylation cancer biomarkers: translation to the clinic. Front. Genet. 10, 1150. 10.3389/fgene.2019.01150.

6. Laird, P.W. (2003). The power and the promise of DNA methylation markers. Nat. Rev. Cancer 3, 253–266. 10.1038/nrc1045.

7. Levy, M.A., McConkey, H., Kerkhof, J., Barat-Houari, M., Bargiacchi, S., Biamino, E., Bralo, M.P., Cappuccio, G., Ciolfi, A., Clarke, A., et al. (2022). Novel diagnostic DNA methylation episignatures expand and refine the epigenetic landscapes of Mendelian disorders. HGG Adv. 3, 100075. 10.1016/j.xhgg.2021.100075.

8. Rots, D., Chater-Diehl, E., Dingemans, A.J.M., Goodman, S.J., Siu, M.T., Cytrynbaum, C., Choufani, S., Hoang, N., Walker, S., Awamleh, Z., et al. (2021). Truncating SRCAP variants outside the Floating-Harbor syndrome locus cause a distinct neurodevelopmental disorder with a specific DNA methylation signature. Am. J. Hum. Genet. 108, 1053–1068. 10.1016/j.ajhg.2021.04.008.

9. Sadikovic, B., Levy, M.A., Kerkhof, J., Aref-Eshghi, E., Schenkel, L., Stuart, A., McConkey, H., Henneman, P., Venema, A., Schwartz, C.E., et al. (2021). Clinical epigenomics: genome-wide DNA methylation analysis for the diagnosis of Mendelian disorders. Genet. Med. 23, 1065–1074. 10.1038/s41436-020-01096-4.

10. Capper, D., Jones, D.T.W., Sill, M., Hovestadt, V., Schrimpf, D., Sturm, D., Koelsche, C., Sahm, F., Chavez, L., Reuss, D.E., et al. (2018). DNA methylation-based classification of central nervous system tumours. Nature 555, 469–474. 10.1038/nature26000.

11. Kerachian, M.A., Azghandi, M., Mozaffari-Jovin, S., and Thierry, A.R. (2021). Guidelines for pre-analytical conditions for assessing the methylation of circulating cell-free DNA. Clin. Epigenetics 13, 193. 10.1186/s13148-021-01182-7.

12. Chen, J., Gatev, E., Everson, T., Conneely, K.N., Koen, N., Epstein, M.P., Kobor, M.S., Zar, H.J., Stein, D.J., and Hüls, A. (2023). Pruning and thresholding approach for methylation risk scores in multi-ancestry populations. Epigenetics 18, 2187172. 10.1080/15592294.2023.2187172.

13. Mannens, M.M.A.M., Lombardi, M.P., Alders, M., Henneman, P., and Bliek, J. (2022). Further introduction of DNA methylation (dnam) arrays in regular diagnostics. Front. Genet. 13, 831452. 10.3389/fgene.2022.831452.

14. Zeng, Y., Jain, R., Lam, M., Ahmed, M., Guo, H., Xu, W., Zhong, Y., Wei, G.-H., Xu, W., and He, H.H. (2023). DNA methylation modulated genetic variant effect on gene transcriptional regulation. Genome Biol. 24, 285. 10.1186/s13059-023-03130-5.

15. Kaluscha, S., Domcke, S., Wirbelauer, C., Stadler, M.B., Durdu, S., Burger, L., and Schübeler, D. (2022). Evidence that direct inhibition of transcription factor binding is the prevailing mode of gene and repeat repression by DNA methylation. Nat. Genet. 54, 1895–1906. 10.1038/s41588-022-01241-6.

16. Luo, Q., Dwaraka, V.B., Chen, Q., Tong, H., Zhu, T., Seale, K., Raffaele, J.M., Zheng, S.C., Mendez, T.L., Chen, Y., et al. (2023). A meta-analysis of immune-cell fractions at high resolution reveals novel associations with common phenotypes and health outcomes. Genome Med. 15, 59. 10.1186/s13073-023-01211-5.

17. Campagna, M.P., Xavier, A., Lechner-Scott, J., Maltby, V., Scott, R.J., Butzkueven, H., Jokubaitis, V.G., and Lea, R.A. (2021). Epigenome-wide association studies: current knowledge, strategies and recommendations. Clin. Epigenetics 13, 214. 10.1186/s13148-021-01200-8.

18. Rakyan, V.K., Down, T.A., Balding, D.J., and Beck, S. (2011). Epigenome-wide association studies for common human diseases. Nat. Rev. Genet. 12, 529–541. 10.1038/nrg3000.

19. Laird, P.W. (2010). Principles and challenges of genomewide DNA methylation analysis. Nat. Rev. Genet. 11, 191–203. 10.1038/nrg2732.

20. Iqbal, W., and Zhou, W. (2023). Computational Methods for Single-cell DNA Methylome Analysis. Genomics Proteomics Bioinformatics 21, 48–66. 10.1016/j.gpb.2022.05.007.

21. Karemaker, I.D., and Vermeulen, M. (2018). Single-Cell DNA Methylation Profiling: Technologies and Biological Applications. Trends Biotechnol. 36, 952–965. 10.1016/j.tibtech.2018.04.002.

22. Tian, W., Zhou, J., Bartlett, A., Zeng, Q., Liu, H., Castanon, R.G., Kenworthy, M., Altshul, J., Valadon, C., Aldridge, A., et al. (2023). Single-cell DNA methylation and 3D genome architecture in the human brain. Science 382, eadf5357. 10.1126/science.adf5357.

23. Zhou, W., Dinh, H.Q., Ramjan, Z., Weisenberger, D.J., Nicolet, C.M., Shen, H., Laird, P.W., and Berman, B.P. (2018). DNA methylation loss in late-replicating domains is linked to mitotic cell division. Nat. Genet. 50, 591–602. 10.1038/s41588-018-0073-4.

24. Loyfer, N., Magenheim, J., Peretz, A., Cann, G., Bredno, J., Klochendler, A., Fox-Fisher, I., Shabi-Porat, S., Hecht, M., Pelet, T., et al. (2023). A DNA methylation atlas of normal human cell types. Nature 613, 355–364. 10.1038/s41586-022-05580-6.

25. Meissner, A., Gnirke, A., Bell, G.W., Ramsahoye, B., Lander, E.S., and Jaenisch, R. (2005). Reduced representation bisulfite sequencing for comparative high-resolution DNA methylation analysis. Nucleic Acids Res. 33, 5868–5877. 10.1093/nar/gki901.

26. Vermeulen, C., Pagès-Gallego, M., Kester, L., Kranendonk, M.E.G., Wesseling, P., Verburg, N., de Witt Hamer, P., Kooi, E.J., Dankmeijer, L., van der Lugt, J., et al. (2023). Ultra-fast deep-learned CNS tumour classification during surgery. Nature 622, 842–849. 10.1038/s41586-023-06615-2.

27. Bibikova, M., Barnes, B., Tsan, C., Ho, V., Klotzle, B., Le, J.M., Delano, D., Zhang, L., Schroth, G.P., Gunderson, K.L., et al. (2011). High density DNA methylation array with single CpG site resolution. Genomics 98, 288–295. 10.1016/j.ygeno.2011.07.007.

28. Maden, S.K., Thompson, R.F., Hansen, K.D., and Nellore, A. (2021). Human methylome variation across Infinium 450K data on the Gene Expression Omnibus. NAR Genom. Bioinform. 3, lqab025. 10.1093/nargab/lqab025.

29. Hawe, J.S., Wilson, R., Schmid, K.T., Zhou, L., Lakshmanan, L.N., Lehne, B.C., Kühnel, B., Scott, W.R., Wielscher, M., Yew, Y.W., et al. (2022). Genetic variation influencing DNA methylation provides insights into molecular mechanisms regulating genomic function. Nat. Genet. 54, 18–29. 10.1038/s41588-021-00969-x.

30. Min, J.L., Hemani, G., Hannon, E., Dekkers, K.F., Castillo-Fernandez, J., Luijk, R., Carnero-Montoro, E., Lawson, D.J., Burrows, K., Suderman, M., et al. (2021). Genomic and phenotypic insights from an atlas of genetic effects on DNA methylation. Nat. Genet. 53, 1311–1321. 10.1038/s41588-021-00923-x.

31. Thompson, M., Hill, B.L., Rakocz, N., Chiang, J.N., Geschwind, D., Sankararaman, S., Hofer, I., Cannesson, M., Zaitlen, N., and Halperin, E. (2022). Methylation risk scores are associated with a collection of phenotypes within electronic health record systems. NPJ Genom. Med. 7, 50. 10.1038/s41525-022-00320-1.

32. Aref-Eshghi, E., Kerkhof, J., Pedro, V.P., Groupe DI France, Barat-Houari, M., Ruiz-Pallares, N., Andrau, J.-C., Lacombe, D., Van-Gils, J., Fergelot, P., et al. (2020). Evaluation of DNA methylation episignatures for diagnosis and phenotype correlations in 42 mendelian neurodevelopmental disorders. Am. J. Hum. Genet. 106, 356–370. 10.1016/j.ajhg.2020.01.019.

33. Battram, T., Yousefi, P., Crawford, G., Prince, C., Sheikhali Babaei, M., Sharp, G., Hatcher, C., Vega-Salas, M.J., Khodabakhsh, S., Whitehurst, O., et al. (2022). The EWAS Catalog: a database of epigenome-wide association studies. Wellcome Open Res. 7, 41. 10.12688/wellcomeopenres.17598.2.

34. Li, M., Zou, D., Li, Z., Gao, R., Sang, J., Zhang, Y., Li, R., Xia, L., Zhang, T., Niu, G., et al. (2019). EWAS Atlas: a curated knowledgebase of epigenome-wide association studies. Nucleic Acids Res. 47, D983–D988. 10.1093/nar/gky1027.

35. Haghani, A., Li, C.Z., Robeck, T.R., Zhang, J., Lu, A.T., Ablaeva, J., Acosta-Rodríguez, V.A., Adams, D.M., Alagaili, A.N., Almunia, J., et al. (2023). DNA methylation networks underlying mammalian traits. Science 381, eabq5693. 10.1126/science.abq5693.

36. Ding, W., Kaur, D., Horvath, S., and Zhou, W. (2023). Comparative epigenome analysis using Infinium DNA methylation BeadChips. Brief. Bioinformatics 24. 10.1093/bib/bbac617.

37. Arneson, A., Haghani, A., Thompson, M.J., Pellegrini, M., Kwon, S.B., Vu, H., Maciejewski, E., Yao, M., Li, C.Z., Lu, A.T., et al. (2022). A mammalian methylation array for profiling methylation levels at conserved sequences. Nat. Commun. 13, 783. 10.1038/s41467-022-28355-z.

38. He, B., Zhang, C., Zhang, X., Fan, Y., Zeng, H., Liu, J., Meng, H., Bai, D., Peng, J., Zhang, Q., et al. (2021). Tissue-specific 5-hydroxymethylcytosine landscape of the human genome. Nat. Commun. 12, 4249. 10.1038/s41467-021-24425-w.

39. Cui, X.-L., Nie, J., Ku, J., Dougherty, U., West-Szymanski, D.C., Collin, F., Ellison, C.K., Sieh, L., Ning, Y., Deng, Z., et al. (2020). A human tissue map of 5-hydroxymethylcytosines exhibits tissue specificity through gene and enhancer modulation. Nat. Commun. 11, 6161. 10.1038/s41467-020-20001-w.

40. Bai, D., Zhang, X., Xiang, H., Guo, Z., Zhu, C., and Yi, C. (2024). Simultaneous single-cell analysis of 5mC and 5hmC with SIMPLE-seq. Nat. Biotechnol. 10.1038/s41587-024-02148-9.

41. Wen, L., and Tang, F. (2014). Genomic distribution and possible functions of DNA hydroxymethylation in the brain. Genomics 104, 341–346. 10.1016/j.ygeno.2014.08.020.

42. Li, W., and Liu, M. (2011). Distribution of 5-hydroxymethylcytosine in different human tissues. J. Nucleic Acids 2011, 870726. 10.4061/2011/870726.

43. Zhou, W., Laird, P.W., and Shen, H. (2017). Comprehensive characterization, annotation and innovative use of Infinium DNA methylation BeadChip probes. Nucleic Acids Res. 45, e22. 10.1093/nar/gkw967.

44. Kaur, D., Lee, S.M., Goldberg, D., Spix, N.J., Hinoue, T., Li, H.-T., Dwaraka, V.B., Smith, R., Shen, H., Liang, G., et al. (2023). Comprehensive evaluation of the infinium human methylationepic v2 beadchip. Epigenetics Commun. 3. 10.1186/s43682-023-00021-5.

45. Horvath, S., and Raj, K. (2018). DNA methylation-based biomarkers and the epigenetic clock theory of ageing. Nat. Rev. Genet. 19, 371–384. 10.1038/s41576-018-0004-3.

46. Weidner, C.I., Lin, Q., Koch, C.M., Eisele, L., Beier, F., Ziegler, P., Bauerschlag, D.O., Jöckel, K.-H., Erbel, R., Mühleisen, T.W., et al. (2014). Aging of blood can be tracked by DNA methylation changes at just three CpG sites. Genome Biol. 15, R24. 10.1186/gb-2014-15-2-r24.

47. Zhang, W., Wu, H., and Li, Z. (2021). Complete deconvolution of DNA methylation signals from complex tissues: a geometric approach. Bioinformatics 37, 1052–1059. 10.1093/bioinformatics/btaa930.

48. Xia, D., Leon, A.J., Cabanero, M., Pugh, T.J., Tsao, M.S., Rath, P., Siu, L.L.-Y., Yu, C., Bedard, P.L., Shepherd, F.A., et al. (2020). Minimalist approaches to cancer tissue-of-origin classification by DNA methylation. Mod. Pathol. 33, 1874–1888. 10.1038/s41379-020-0547-7.

49. Luo, C., Liu, H., Xie, F., Armand, E.J., Siletti, K., Bakken, T.E., Fang, R., Doyle, W.I., Stuart, T., Hodge, R.D., et al. (2022). Single nucleus multi-omics identifies human cortical cell regulatory genome diversity. Cell Genomics 2. 10.1016/j.xgen.2022.100107.

50. Lee, D.-S., Luo, C., Zhou, J., Chandran, S., Rivkin, A., Bartlett, A., Nery, J.R., Fitzpatrick, C., O’Connor, C., Dixon, J.R., et al. (2019). Simultaneous profiling of 3D genome structure and DNA methylation in single human cells. Nat. Methods 16, 999–1006. 10.1038/s41592-019-0547-z.

51. Luo, C., Keown, C.L., Kurihara, L., Zhou, J., He, Y., Li, J., Castanon, R., Lucero, J., Nery, J.R., Sandoval, J.P., et al. (2017). Single-cell methylomes identify neuronal subtypes and regulatory elements in mammalian cortex. Science 357, 600–604. 10.1126/science.aan3351.

52. Martens, J.H.A., and Stunnenberg, H.G. (2013). BLUEPRINT: mapping human blood cell epigenomes. Haematologica 98, 1487–1489. 10.3324/haematol.2013.094243.

53. Vu, H., and Ernst, J. (2022). Universal annotation of the human genome through integration of over a thousand epigenomic datasets. Genome Biol. 23, 9. 10.1186/s13059-021-02572-z.

54. ENCODE Project Consortium, Moore, J.E., Purcaro, M.J., Pratt, H.E., Epstein, C.B., Shoresh, N., Adrian, J., Kawli, T., Davis, C.A., Dobin, A., et al. (2022). Author Correction: Expanded encyclopaedias of DNA elements in the human and mouse genomes. Nature 605, E3. 10.1038/s41586-021-04226-3.

55. Zhou, W., Hinoue, T., Barnes, B., Mitchell, O., Iqbal, W., Lee, S.M., Foy, K.K., Lee, K.-H., Moyer, E.J., VanderArk, A., et al. (2022). DNA methylation dynamics and dysregulation delineated by high-throughput profiling in the mouse. Cell Genomics 2. 10.1016/j.xgen.2022.100144.

56. Zhou, W., Triche, T.J., Laird, P.W., and Shen, H. (2018). SeSAMe: reducing artifactual detection of DNA methylation by Infinium BeadChips in genomic deletions. Nucleic Acids Res. 46, e123. 10.1093/nar/gky691.

57. Lee, S.M., Loo, C.E., Prasasya, R.D., Bartolomei, M.S., Kohli, R.M., and Zhou, W. (2024). Low-input and single-cell methods for Infinium DNA methylation BeadChips. Nucleic Acids Res. 52, e38. 10.1093/nar/gkae127.

58. Vaisvila, R., Ponnaluri, V.K.C., Sun, Z., Langhorst, B.W., Saleh, L., Guan, S., Dai, N., Campbell, M.A., Sexton, B.S., Marks, K., et al. (2021). Enzymatic methyl sequencing detects DNA methylation at single-base resolution from picograms of DNA. Genome Res. 31, 1280–1289. 10.1101/gr.266551.120.

59. Groarke, E.M., and Young, N.S. (2019). Aging and Hematopoiesis. Clin. Geriatr. Med. 35, 285–293. 10.1016/j.cger.2019.03.001.

60. Thapa, P., and Farber, D.L. (2019). The role of the thymus in the immune response. Thorac. Surg. Clin. 29, 123–131. 10.1016/j.thorsurg.2018.12.001.

61. Muraro, M.J., Dharmadhikari, G., Grün, D., Groen, N., Dielen, T., Jansen, E., van Gurp, L., Engelse, M.A., Carlotti, F., de Koning, E.J.P., et al. (2016). A Single-Cell Transcriptome Atlas of the Human Pancreas. Cell Syst. 3, 385–394.e3. 10.1016/j.cels.2016.09.002.

62. O’Brien, L.L., Guo, Q., Bahrami-Samani, E., Park, J.-S., Hasso, S.M., Lee, Y.-J., Fang, A., Kim, A.D., Guo, J., Hong, T.M., et al. (2018). Transcriptional regulatory control of mammalian nephron progenitors revealed by multi-factor cistromic analysis and genetic studies. PLoS Genet. 14, e1007181. 10.1371/journal.pgen.1007181.

63. Coskun, M., Troelsen, J.T., and Nielsen, O.H. (2011). The role of CDX2 in intestinal homeostasis and inflammation. Biochim. Biophys. Acta 1812, 283–289. 10.1016/j.bbadis.2010.11.008.

64. Teschendorff, A.E. (2020). A comparison of epigenetic mitotic-like clocks for cancer risk prediction. Genome Med. 12, 56. 10.1186/s13073-020-00752-3.

65. Sender, R., and Milo, R. (2021). The distribution of cellular turnover in the human body. Nat. Med. 27, 45–48. 10.1038/s41591-020-01182-9.

66. Huang, Y., Pastor, W.A., Shen, Y., Tahiliani, M., Liu, D.R., and Rao, A. (2010). The behaviour of 5-hydroxymethylcytosine in bisulfite sequencing. PLoS ONE 5, e8888. 10.1371/journal.pone.0008888.

67. Schutsky, E.K., DeNizio, J.E., Hu, P., Liu, M.Y., Nabel, C.S., Fabyanic, E.B., Hwang, Y., Bushman, F.D., Wu, H., and Kohli, R.M. (2018). Nondestructive, base-resolution sequencing of 5-hydroxymethylcytosine using a DNA deaminase. Nat. Biotechnol. 36, 1083–1090. 10.1038/nbt.4204.

68. Fagerberg, L., Hallström, B.M., Oksvold, P., Kampf, C., Djureinovic, D., Odeberg, J., Habuka, M., Tahmasebpoor, S., Danielsson, A., Edlund, K., et al. (2014). Analysis of the human tissue-specific expression by genome-wide integration of transcriptomics and antibody-based proteomics. Mol. Cell. Proteomics 13, 397–406. 10.1074/mcp.M113.035600.

69. Yamamura, Y., Furuichi, K., Murakawa, Y., Hirabayashi, S., Yoshihara, M., Sako, K., Kitajima, S., Toyama, T., Iwata, Y., Sakai, N., et al. (2021). Identification of candidate PAX2-regulated genes implicated in human kidney development. Sci. Rep. 11, 9123. 10.1038/s41598-021-88743-1.

70. Tsunoda, T., Kakinuma, S., Miyoshi, M., Kamiya, A., Kaneko, S., Sato, A., Tsuchiya, J., Nitta, S., Kawai-Kitahata, F., Murakawa, M., et al. (2019). Loss of fibrocystin promotes interleukin-8-dependent proliferation and CTGF production of biliary epithelium. J. Hepatol. 71, 143–152. 10.1016/j.jhep.2019.02.024.

71. Rocamora-Reverte, L., Melzer, F.L., Würzner, R., and Weinberger, B. (2020). The complex role of regulatory T cells in immunity and aging. Front. Immunol. 11, 616949. 10.3389/fimmu.2020.616949.

72. Horvath, S. (2013). DNA methylation age of human tissues and cell types. Genome Biol. 14, R115. 10.1186/gb-2013-14-10-r115.

73. Hannum, G., Guinney, J., Zhao, L., Zhang, L., Hughes, G., Sadda, S., Klotzle, B., Bibikova, M., Fan, J.-B., Gao, Y., et al. (2013). Genome-wide methylation profiles reveal quantitative views of human aging rates. Mol. Cell 49, 359–367. 10.1016/j.molcel.2012.10.016.

74. Moss, J., Magenheim, J., Neiman, D., Zemmour, H., Loyfer, N., Korach, A., Samet, Y., Maoz, M., Druid, H., Arner, P., et al. (2018). Comprehensive human cell-type methylation atlas reveals origins of circulating cell-free DNA in health and disease. Nat. Commun. 9, 5068. 10.1038/s41467-018-07466-6.

75. Guintivano, J., Arad, M., Gould, T.D., Payne, J.L., and Kaminsky, Z.A. (2014). Antenatal prediction of postpartum depression with blood DNA methylation biomarkers. Mol. Psychiatry 19, 560–567. 10.1038/mp.2013.62.

76. Zhang, Z., Lee, M.K., Perreard, L., Kelsey, K.T., Christensen, B.C., and Salas, L.A. (2022). Navigating the hydroxymethylome: experimental biases and quality control tools for the tandem bisulfite and oxidative bisulfite Illumina microarrays. Epigenomics 14, 139–152. 10.2217/epi-2021-0490.

77. Tiedemann, R.L., Eden, H.E., Huang, Z., Robertson, K.D., and Rothbart, S.B. (2021). Distinguishing active versus passive DNA demethylation using illumina methylationepic beadchip microarrays. Methods Mol. Biol. 2272, 97–140. 10.1007/978-1-0716-1294-1_7.

78. Zhou, W., Johnson, B.K., Morrison, J., Beddows, I., Eapen, J., Katsman, E., Semwal, A., Habib, W.A., Heo, L., Laird, P.W., et al. (2024). BISCUIT: an efficient, standards-compliant tool suite for simultaneous genetic and epigenetic inference in bulk and single-cell studies. Nucleic Acids Res. 52, e32. 10.1093/nar/gkae097.

79. Sherry, S.T., Ward, M.H., Kholodov, M., Baker, J., Phan, L., Smigielski, E.M., and Sirotkin, K. (2001). dbSNP: the NCBI database of genetic variation. Nucleic Acids Res. 29, 308–311. 10.1093/nar/29.1.308.

80. ENCODE Project Consortium, Moore, J.E., Purcaro, M.J., Pratt, H.E., Epstein, C.B., Shoresh, N., Adrian, J., Kawli, T., Davis, C.A., Dobin, A., et al. (2020). Expanded encyclopaedias of DNA elements in the human and mouse genomes. Nature 583, 699– 710. 10.1038/s41586-020-2493-4.

81. Gao, G.F., Parker, J.S., Reynolds, S.M., Silva, T.C., Wang, L.-B., Zhou, W., Akbani, R., Bailey, M., Balu, S., Berman, B.P., et al. (2019). Before and after: comparison of legacy and harmonized TCGA genomic data commons’ data. Cell Syst. 9, 24–34.e10. 10.1016/j.cels.2019.06.006.

82. Gunasekara, C.J., MacKay, H., Scott, C.A., Li, S., Laritsky, E., Baker, M.S., Grimm, S.L., Jun, G., Li, Y., Chen, R., et al. (2023). Systemic interindividual epigenetic variation in humans is associated with transposable elements and under strong genetic control. Genome Biol. 24, 2. 10.1186/s13059-022-02827-3.

83. Gunasekara, C.J., Scott, C.A., Laritsky, E., Baker, M.S., MacKay, H., Duryea, J.D., Kessler, N.J., Hellenthal, G., Wood, A.C., Hodges, K.R., et al. (2019). A genomic atlas of systemic interindividual epigenetic variation in humans. Genome Biol. 20, 105. 10.1186/s13059-019-1708-1.

84. Skaar, D.A., Li, Y., Bernal, A.J., Hoyo, C., Murphy, S.K., and Jirtle, R.L. (2012). The human imprintome: regulatory mechanisms, methods of ascertainment, and roles in disease susceptibility. ILAR J. 53, 341–358. 10.1093/ilar.53.3-4.341.

85. Wang, T., Luo, M., Berrios, K.N., Schutsky, E.K., Wu, H., and Kohli, R.M. (2021). Bisulfite-Free Sequencing of 5-Hydroxymethylcytosine with APOBEC-Coupled Epigenetic Sequencing (ACE-Seq). Methods Mol. Biol. 2198, 349–367. 10.1007/978-1-0716-0876-0_27.

86. Subramanian, A., Tamayo, P., Mootha, V.K., Mukherjee, S., Ebert, B.L., Gillette, M.A., Paulovich, A., Pomeroy, S.L., Golub, T.R., Lander, E.S., et al. (2005). Gene set enrichment analysis: a knowledge-based approach for interpreting genome-wide expression profiles. Proc Natl Acad Sci USA 102, 15545–15550. 10.1073/pnas.0506580102.

87. Xie, Z., Bailey, A., Kuleshov, M.V., Clarke, D.J.B., Evangelista, J.E., Jenkins, S.L., Lachmann, A., Wojciechowicz, M.L., Kropiwnicki, E., Jagodnik, K.M., et al. (2021). Gene Set Knowledge Discovery with Enrichr. Curr. Protoc. 1, e90. 10.1002/cpz1.90.

88. Teschendorff, A.E., Breeze, C.E., Zheng, S.C., and Beck, S. (2017). A comparison of reference-based algorithms for correcting cell-type heterogeneity in Epigenome-Wide Association Studies. BMC Bioinformatics 18, 105. 10.1186/s12859-017-1511-5.

89. Hammal, F., de Langen, P., Bergon, A., Lopez, F., and Ballester, B. (2022). ReMap 2022: a database of Human, Mouse, Drosophila and Arabidopsis regulatory regions from an integrative analysis of DNA-binding sequencing experiments. Nucleic Acids Res. 50, D316–D325. 10.1093/nar/gkab996.

90. Su, A.I., Wiltshire, T., Batalov, S., Lapp, H., Ching, K.A., Block, D., Zhang, J., Soden, R., Hayakawa, M., Kreiman, G., et al. (2004). A gene atlas of the mouse and human protein-encoding transcriptomes. Proc Natl Acad Sci USA 101, 6062–6067. 10.1073/pnas.0400782101.

91. Kuleshov, M.V., Jones, M.R., Rouillard, A.D., Fernandez, N.F., Duan, Q., Wang, Z., Koplev, S., Jenkins, S.L., Jagodnik, K.M., Lachmann, A., et al. (2016). Enrichr: a comprehensive gene set enrichment analysis web server 2016 update. Nucleic Acids Res. 44, W90–7. 10.1093/nar/gkw377.

92. Chen, E.Y., Tan, C.M., Kou, Y., Duan, Q., Wang, Z., Meirelles, G.V., Clark, N.R., and Ma’ayan, A. (2013). Enrichr: interactive and collaborative HTML5 gene list enrichment analysis tool. BMC Bioinformatics 14, 128. 10.1186/1471-2105-14-128.

93. Pelegí-Sisó, D., de Prado, P., Ronkainen, J., Bustamante, M., and González, J.R. (2021). methylclock: a Bioconductor package to estimate DNA methylation age. Bioinformatics 37, 1759–1760. 10.1093/bioinformatics/btaa825.

94. Sollis, E., Mosaku, A., Abid, A., Buniello, A., Cerezo, M., Gil, L., Groza, T., Güneş, O., Hall, P., Hayhurst, J., et al. (2023). The NHGRI-EBI GWAS Catalog: knowledgebase and deposition resource. Nucleic Acids Res. 51, D977–D985. 10.1093/nar/gkac1010.

