## Supplemental Figures for "Scalable Screening of Ternary-Code DNA Methylation Dynamics Associated with Human Traits"

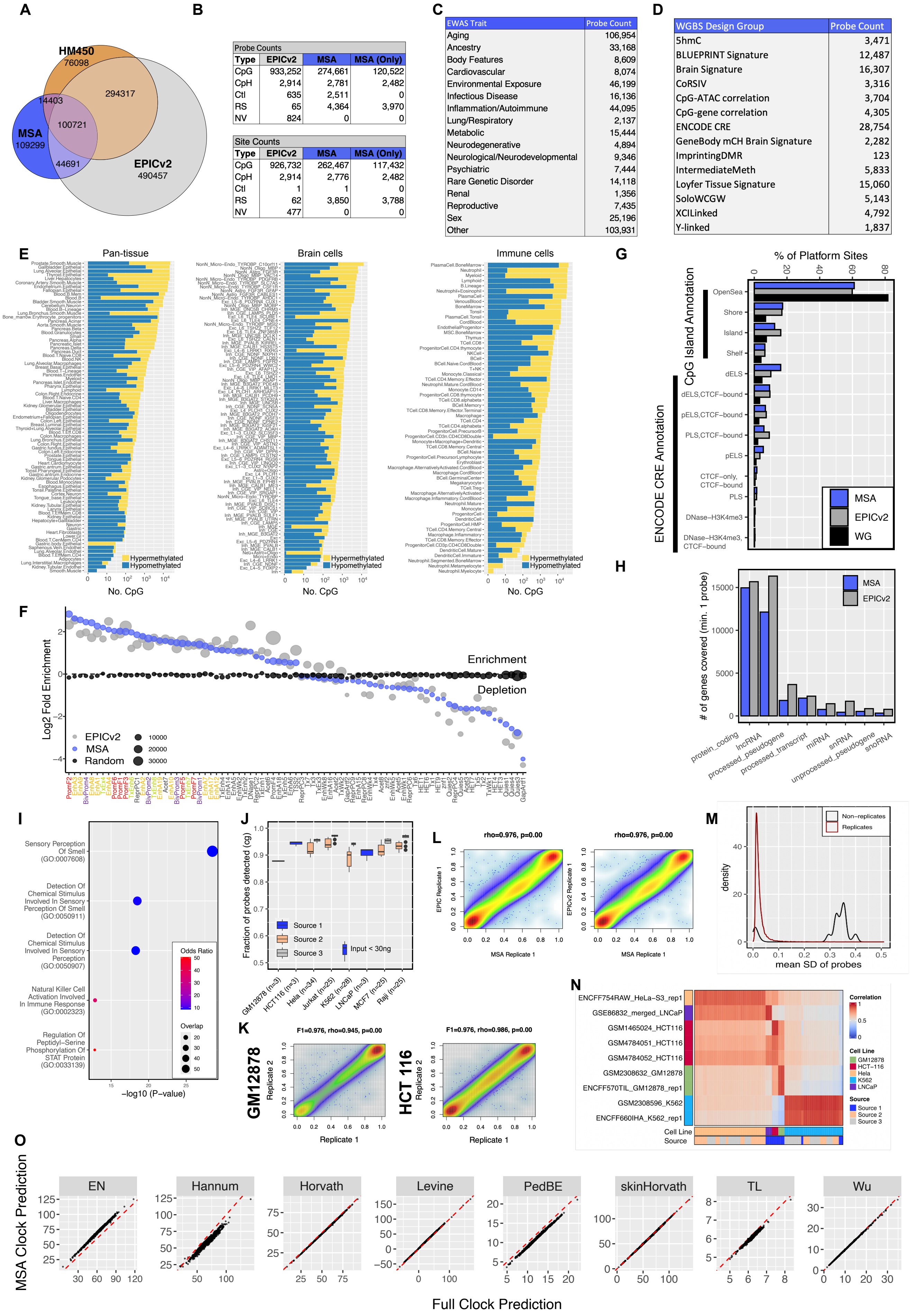


| Probe Counts | | | |
|---|---|---|---|
| Type | EPICv2 | MSA | MSA (Only) |
| CpG | 933,252 | 274,661 | 120,522 |
| CpH | 2,914 | 2,781 | 2,482 |
| Ctl | 635 | 2,511 | 0 |
| RS | 65 | 4,364 | 3,970 |
| NV | 824 | 0 | 0 |

| Site Counts | | | |
|---|---|---|---|
| Type | EPICv2 | MSA | MSA (Only) |
| CpG | 926,732 | 262,467 | 117,432 |
| CpH | 2,914 | 2,776 | 2,482 |
| Ctl | 1 | 1 | 0 |
| RS | 62 | 3,850 | 3,788 |
| NV | 477 | 0 | 0 |

| EWAS Trait | Probe Count |
|---|---|
| Aging | 106,954 |
| Ancestry | 33,168 |
| Body Features | 8,609 |
| Cardiovascular | 8,074 |
| Environmental Exposure | 46,199 |
| Infectious Disease | 16,136 |
| Inflammation/Autoimmune | 44,095 |
| Lung/Respiratory | 2,137 |
| Metabolic | 15,444 |
| Neurodegenerative | 4,894 |
| Neurological/Neurodevelopmental | 9,346 |
| Psychiatric | 7,444 |
| Rare Genetic Disorder | 14,118 |
| Renal | 1,356 |
| Reproductive | 7,435 |
| Sex | 25,196 |
| Other | 103,931 |

| WGBS Design Group | Probe Count |
|---|---|
| 5hmC | 3,471 |
| BLUEPRINT Signature | 12,487 |
| Brain Signature | 16,307 |
| CoRSIV | 3,316 |
| CpG-ATAC correlation | 3,704 |
| CpG-gene correlation | 4,305 |
| ENCODE CRE | 28,754 |
| GeneBody mCH Brain Signature | 2,282 |
| ImprintingDMR | 123 |
| IntermediateMeth | 5,833 |
| Loyfer Tissue Signature | 15,060 |
| SoloWCGW | 5,143 |
| XCILinked | 4,792 |
| Y-linked | 1,837 |

**Figure S1: MSA probe selection summary and technical validation.** (A) Venn diagram displaying the number of genomic sites covered on MSA and the overlap with EPICv2 and HM450 sites. (B) Table of total probe counts for different probe types on EPICv2, MSA, and novel MSA designs not found on previous Infinium platforms. (C) Table of probe counts for major EWAS trait group annotations. Probes are deduplicated within major trait groups. (D) Table showing deduplicated probe counts for whole genome methylation design groups targeted (E) Numbers of hyper and hypo methylated signature designs for cell types on MSA. Counts include one vs. all and subtype contrasts. (F) Enrichment of MSA probes in full stack ChromHMM states compared with EPICv2 and a random selection. (G) Percentage of sites in cis-regulatory elements and CpG islands for MSA and EPICv2 compared to whole genome CpGs. (H) Number of genes covered with a minimum of one probe (within 1500bp of the TSS) for MSA and EPICv2. (I) Gene ontology results for genes not targeted (minimum of one probe within 1500bp of TSS) on MSA. (J) Boxplots showing probe detection rates for different cell lines profiled on MSA. (K) Scatter plots showing beta values for cell line technical replicates. (L) Scatter plots showing beta values for cell line replicates profiled on EPIC (y-axis) and MSA (x-axis) (M) Density plots showing the distributions of within-sample mean standard deviations for replicate probe designs compared to within-sample mean standard deviations for non-replicate designs. (N) Heatmap of beta value correlations between cell line samples profiled on MSA and publicly available WGBS data of related cell lines. (O) Correlation of age estimates using full clocks and MSA probe-only clocks on TCGA normal tissues.

A

| Tissue | Total | Sex (M/F) | 5mC | 5hmC |
|---|---|---|---|---|
| Adrenal | 6 | 3/3 | ✓ | ✓ |
| Bladder | 6 | 3/3 | ✓ | ✓ |
| CD4+ T cells | 3 | 2/1 | ✓ | ✗ |
| CD8+ T cells | 3 | 2/1 | ✓ | ✗ |
| Cerebellum | 2 | 0/2 | ✓ | ✓ |
| Colon | 9 | 4/5 | ✓ | ✓ |
| Esophagus | 6 | 3/3 | ✓ | ✓ |
| Eye | 2 | 2/0 | ✓ | ✓ |
| Heart | 6 | 3/3 | ✓ | ✓ |
| Kidney | 5 | 2/3 | ✓ | ✓ |
| Larynx | 5 | 4/1 | ✓ | ✓ |
| Liver | 4 | 2/2 | ✓ | ✓ |
| Lung | 4 | 3/1 | ✓ | ✓ |
| Lymph Node | 3 | 1/2 | ✓ | ✓ |
| Monocytes | 3 | 2/1 | ✓ | ✗ |
| Motor cortex | 3 | 1/2 | ✓ | ✓ |
| NK cells | 3 | 1/2 | ✓ | ✗ |
| Ovary | 3 | 0/3 | ✓ | ✓ |
| PBMC | 3 | 2/1 | ✓ | ✗ |
| Pancreas | 6 | 4/2 | ✓ | ✓ |
| Placenta | 3 | 0/3 | ✓ | ✓ |
| Prostate | 4 | 4/0 | ✓ | ✓ |
| Skin | 6 | 3/3 | ✓ | ✓ |
| Spinal Cord | 1 | 0/1 | ✓ | ✓ |
| Spleen | 5 | 2/3 | ✓ | ✓ |
| Stomach | 6 | 3/3 | ✓ | ✓ |
| Testis | 6 | 6/0 | ✓ | ✓ |
| Thymus | 6 | 3/3 | ✓ | ✓ |
| Thyroid | 5 | 3/2 | ✓ | ✓ |
| Total T cells | 3 | 2/1 | ✓ | ✗ |
| Trachea | 5 | 3/2 | ✓ | ✓ |

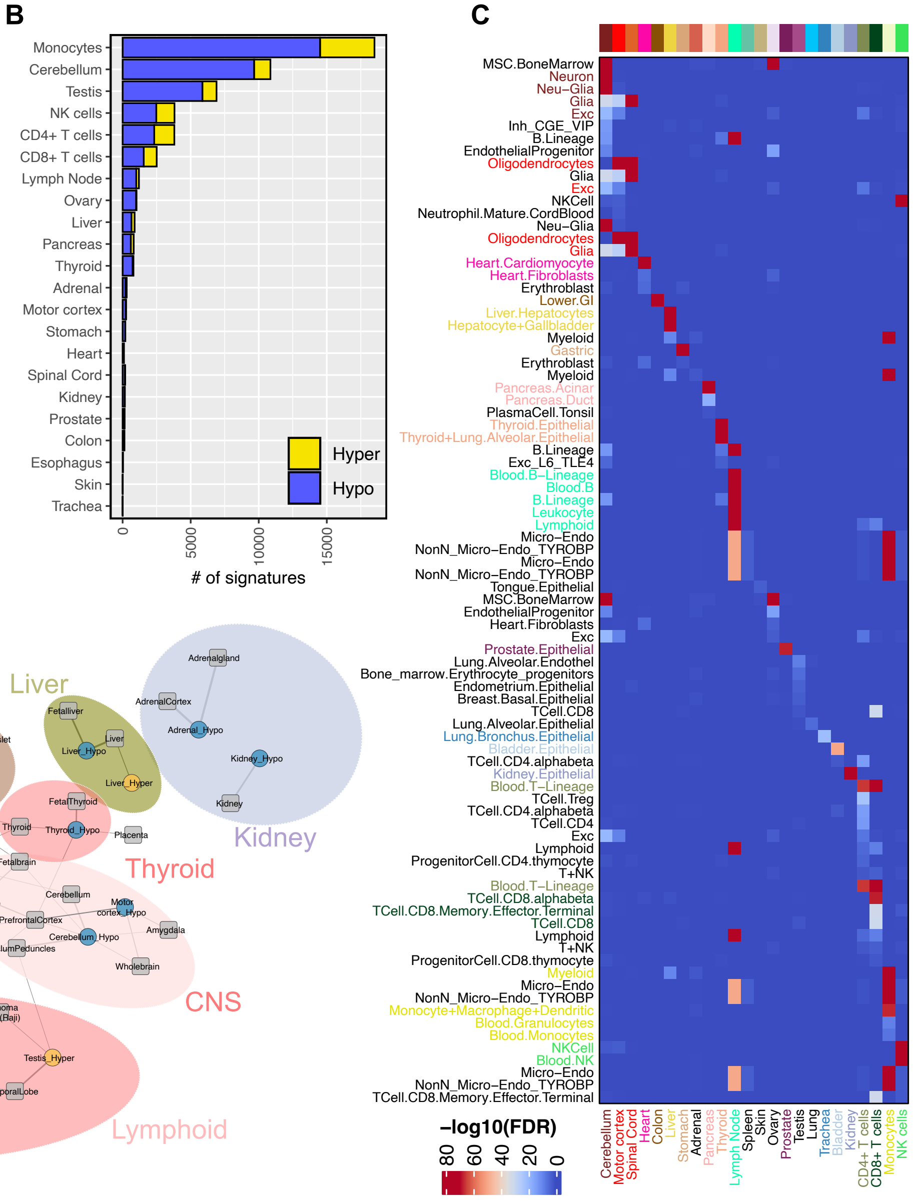


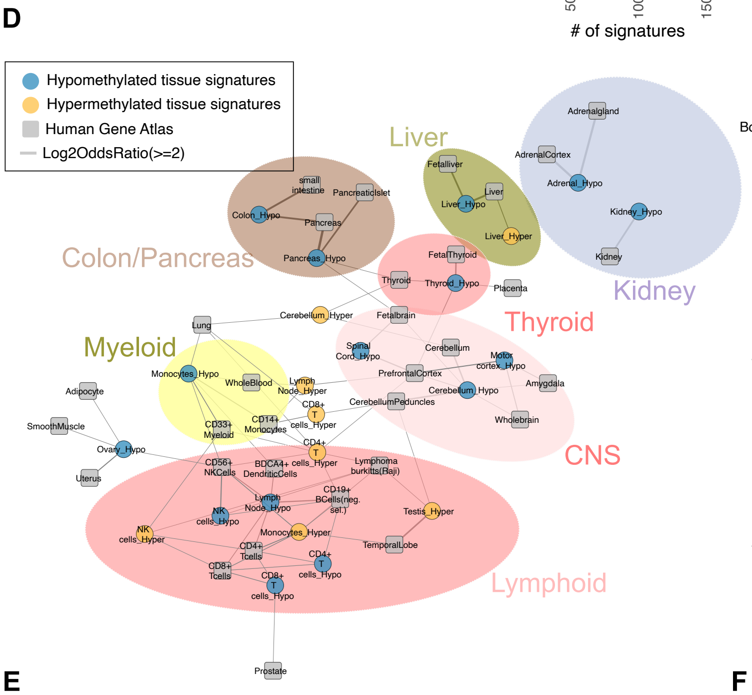


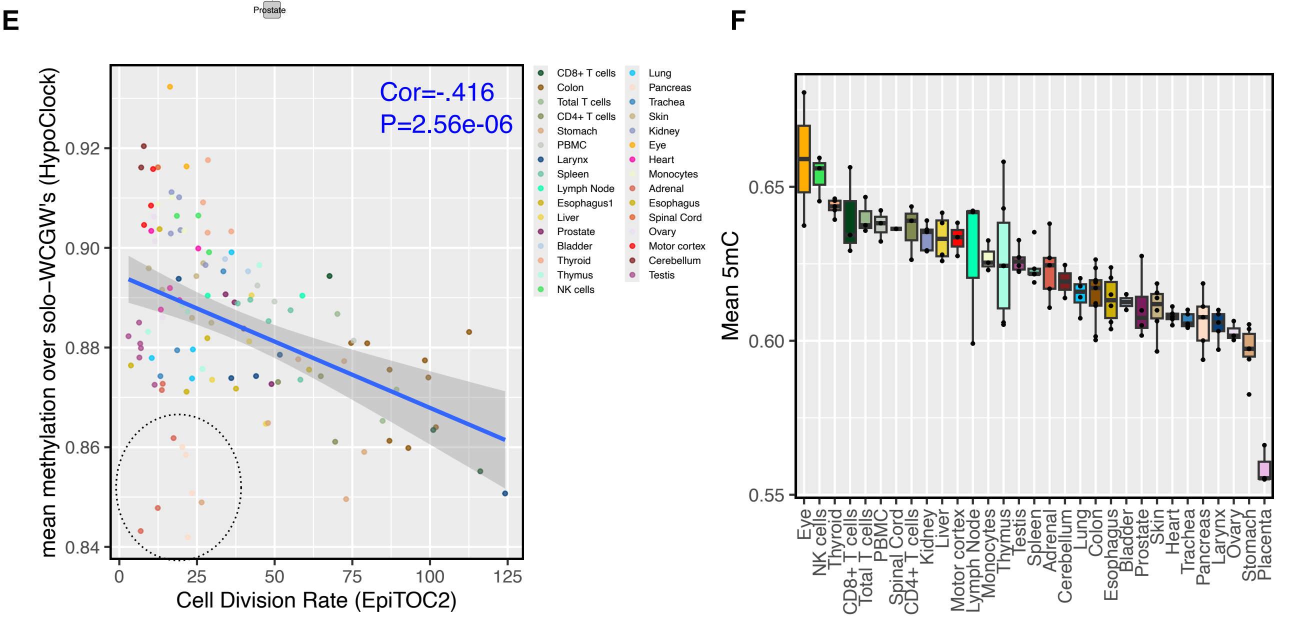

**Figure S2: Tissue-specific methylation detected on MSA.** (A) Table of human tissues profiled on MSA for 5hmC and 5mC in this study (B) Counts of high confidence hyper (yellow) and hypo (blue) methylated CpG markers identified for each tissue type profiled. (C) Heatmap showing enrichment of cell-specific CpGs identified in the current study using MSA (columns) in cell-specific curations from publicly available WGBS data (rows) (D) Network showing gene groups linked to tissue-specific CpGs identified in this study (blue and orange nodes) and tissue-specific genes from the Human Gene Atlas (grey nodes). Edges represent the enrichment of gene sets. (E) Correlation of cell division estimates computed with PRC2 methylation (x-axis) and solo-WCGW methylation (y-axis) for MSA profiled tissues. (F) Mean CpG 5modC levels for tissues profiled with MSA in this study.

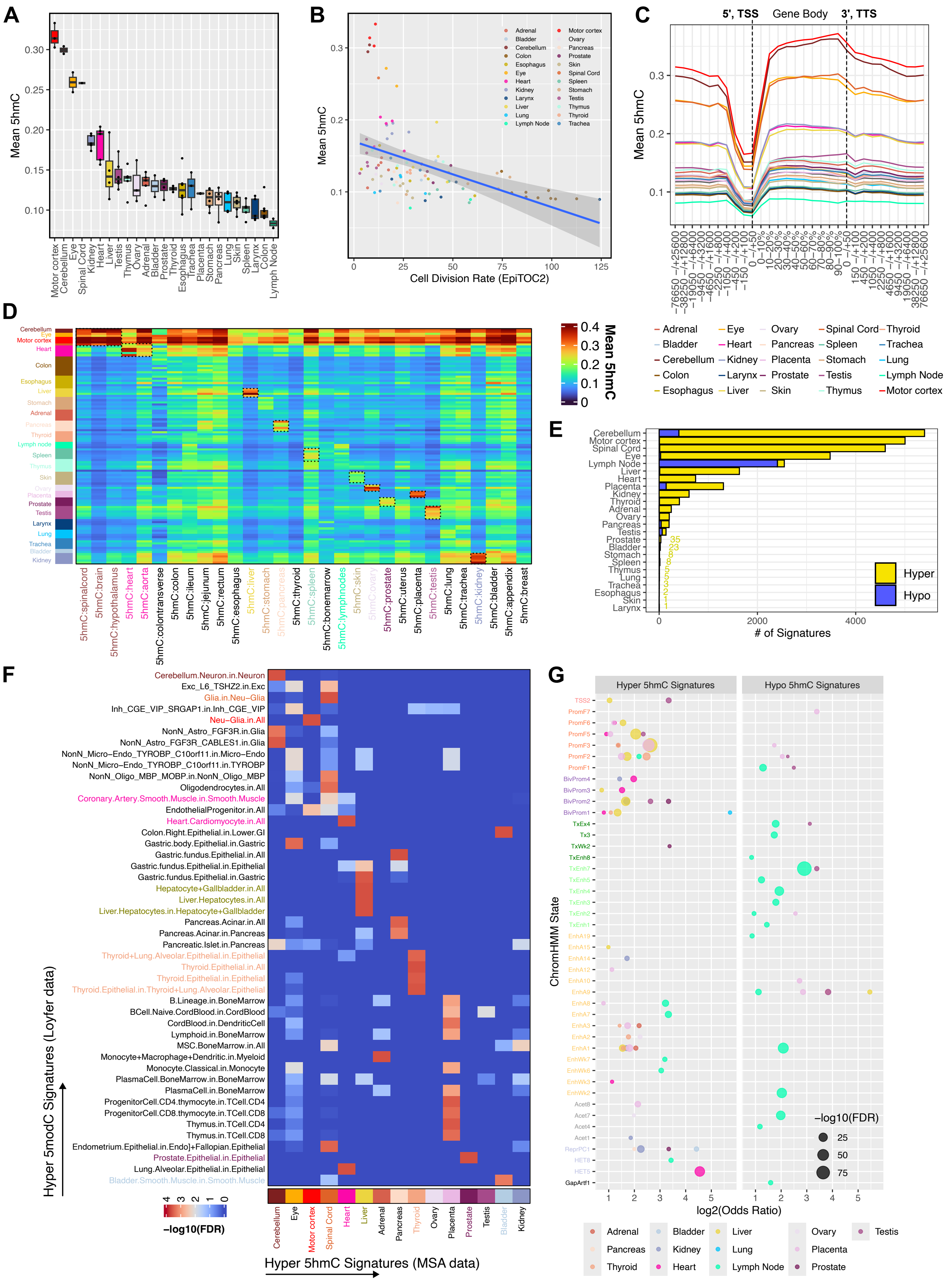
A
Mean 5hmC
Motor cortex
Cerebellum
Eye
Spinal Cord
Kidney
Heart
Liver
Testis
Thymus
Ovary
Adrenal
Bladder
Prostate
Thyroid
Esophagus
Trachea
Placenta
Stomach
Pancreas
Lung
Skin
Spleen
Larynx
Colon
Lymph Node
B
Mean 5hmC
Cell Division Rate (EpiTOC2)
C
5', TSS
Gene Body
3', TTS
Mean 5hmC
D
Mean 5hmC
5hmC:spinalcord
5hmC:brain
5hmC:hypothalamus
5hmC:heart
5hmC:aorta
5hmC:colontransverse
5hmC:colon
5hmC:ileum
5hmC:jejunum
5hmC:rectum
5hmC:esophagus
5hmC:liver
5hmC:stomach
5hmC:pancreas
5hmC:thyroid
5hmC:spleen
5hmC:bonemarrow
5hmC:lymphnodes
5hmC:skin
5hmC:ovary
5hmC:prostate
5hmC:uterus
5hmC:placenta
5hmC:testis
5hmC:lung
5hmC:trachea
5hmC:kidney
5hmC:bladder
5hmC:appendix
5hmC:breast
E
### of Signatures
Hyper
Hypo
F
Hyper 5modC Signatures (Loyfer data)
Hyper 5hmC Signatures (MSA data)
−log10(FDR)
Cerebellum.Neuron.in.Neuron
Exc_L6_TSHZ2.in.Exc
Glia.in.Neu−Glia
Inh_CGE_VIP_SRGAP1.in.Inh_CGE_VIP
Neu−Glia.in.All
NonN_Astro_FGF3R.in.Glia
NonN_Astro_FGF3R_CABLES1.in.Glia
NonN_Micro−Endo_TYROBP_C10orf11.in.Micro−Endo
NonN_Micro−Endo_TYROBP_C10orf11.in.TYROBP
NonN_Oligo_MBP_MOBP.in.NonN_Oligo_MBP
Oligodendrocytes.in.All
Coronary.Artery.Smooth.Muscle.in.Smooth.Muscle
EndothelialProgenitor.in.All
Heart.Cardiomyocyte.in.All
Colon.Right.Epithelial.in.Lower.GI
Gastric.body.Epithelial.in.Gastric
Gastric.fundus.Epithelial.in.All
Gastric.fundus.Epithelial.in.Epithelial
Gastric.fundus.Epithelial.in.Gastric
Hepatocyte+Gallbladder.in.All
Liver.Hepatocytes.in.All
Liver.Hepatocytes.in.Hepatocyte+Gallbladder
Pancreas.Acinar.in.All
Pancreas.Acinar.in.Pancreas
Pancreatic.Islet.in.Pancreas
Thyroid+Lung.Alveolar.Epithelial.in.Epithelial
Thyroid.Epithelial.in.All
Thyroid.Epithelial.in.Epithelial
Thyroid.Epithelial.in.Thyroid+Lung.Alveolar.Epithelial
B.Lineage.in.BoneMarrow
BCell.Naive.CordBlood.in.CordBlood
CordBlood.in.DendriticCell
Lymphoid.in.BoneMarrow
MSC.BoneMarrow.in.All
Monocyte+Macrophage+Dendritic.in.Myeloid
Monocyte.Classical.in.Monocyte
PlasmaCell.BoneMarrow.in.BoneMarrow
PlasmaCell.in.BoneMarrow
ProgenitorCell.CD4.thymocyte.in.TCell.CD4
ProgenitorCell.CD8.thymocyte.in.TCell.CD8
Thymus.in.TCell.CD4
Thymus.in.TCell.CD8
Endometrium.Epithelial.in.Endo]+Fallopian.Epithelial
Prostate.Epithelial.in.Epithelial
Lung.Alveolar.Epithelial.in.Epithelial
Bladder.Smooth.Muscle.in.Smooth.Muscle
G
Hyper 5hmC Signatures
Hypo 5hmC Signatures
ChromHMM State
log2(Odds Ratio)
−log10(FDR)

**Figure S3: MSA detects 5hmC in human tissues.** (A) Boxplot showing mean global 5hmC levels across tissues (B) Correlation of cell division estimates computed with PRC2 methylation (x-axis) with mean global 5hmC levels (C) Meta gene plot showing mean 5hmC levels across all genes relative to the transcription start site (D) Heatmap showing mean 5hmC levels for each sample (rows) across the 5hmC tissue-specific design groups created during array development (columns) (E) Counts of high confidence hyper (yellow) and hypo (blue) 5hmC CpG markers for each tissue type profiled. (F) Heatmap showing the enrichment of 5hmC signatures in tissue-specific hypermodified (5mC + 5hmC) signatures curated during array development from WGBS data (G) Enrichment of hyper and hypo 5hmC tissue-specific CpGs in different full-stack ChromHMM chromatin states (FDR < .05)

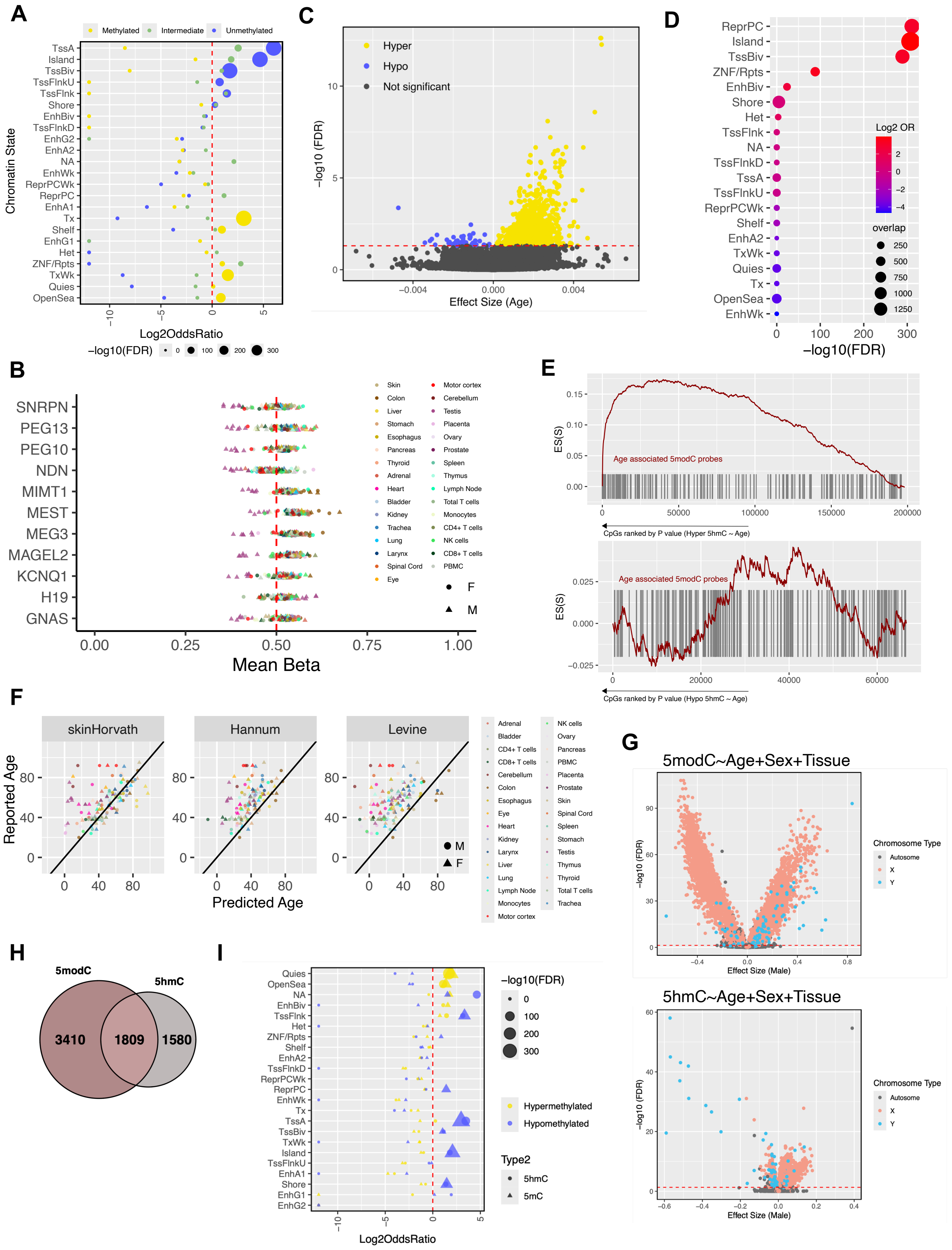


**Figure S4: 5mC and 5hmC methylation biology in imprinting, aging, and sex specificities.** (A) Dotplot showing enrichment of constitutive hypo, hyper, or intermediately methylated CpG classes across consensus ChromHMM states. (B) Mean methylation over intermediately methylated probes linked to known imprinting genes. (C) Volcano plot from aging EWAS (D) Dot plot showing enrichment of age-associated CpGs in consensus ChromHMM states. (E) Correlation of predicted age computed with three epigenetic clocks with actual age for tissues profiled on MSA. (F) Set enrichment analysis showing enrichment of age-associated 5modC probes in the ranked list of age-associated hyper 5hmC probes (top) and age-associated hypo 5hmC (bottom) (G) Volcano plots for Sex EWAS for 5modC (top panel) and 5hmC (bottom panel). (H) Venn diagram showing overlap probes with sex-associated 5hmC and 5modC (I) Dot plot showing enrichment of sex-associated 5modC and 5hmC across consensus ChromHMM states.

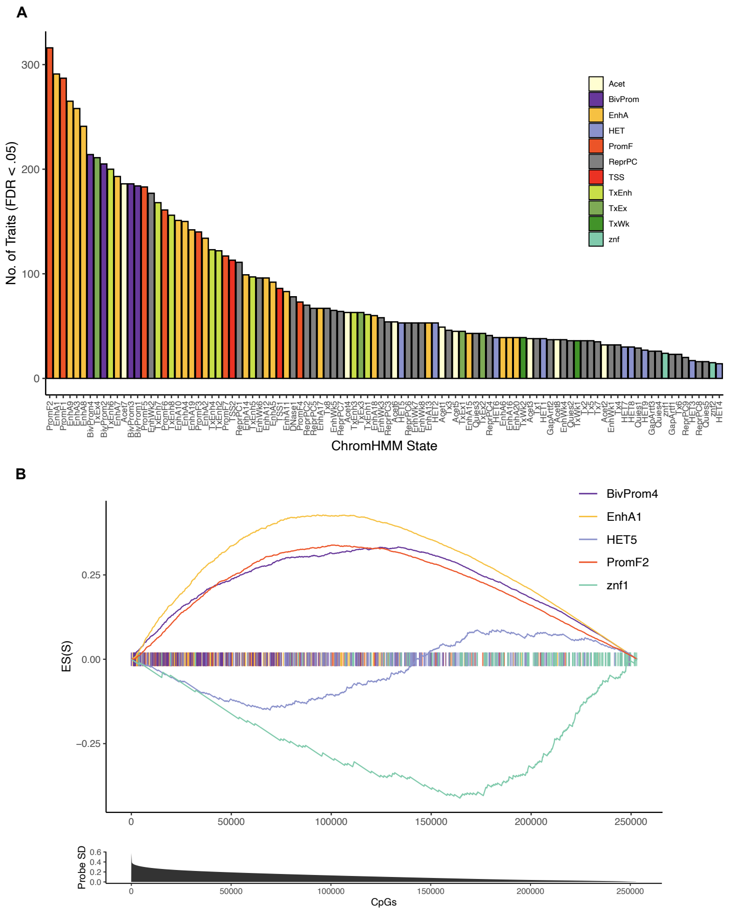


**Figure S5: Tissue and chromatin context of human trait associations.** (A) The number of EWAS traits significantly enriched for each full-stack ChromHMM chromatin state. (B) Set enrichment plot showing where EWAS hits from the selected chromatin states are enriched in the ranked list of most variable probes in MSA-profiled human tissues.

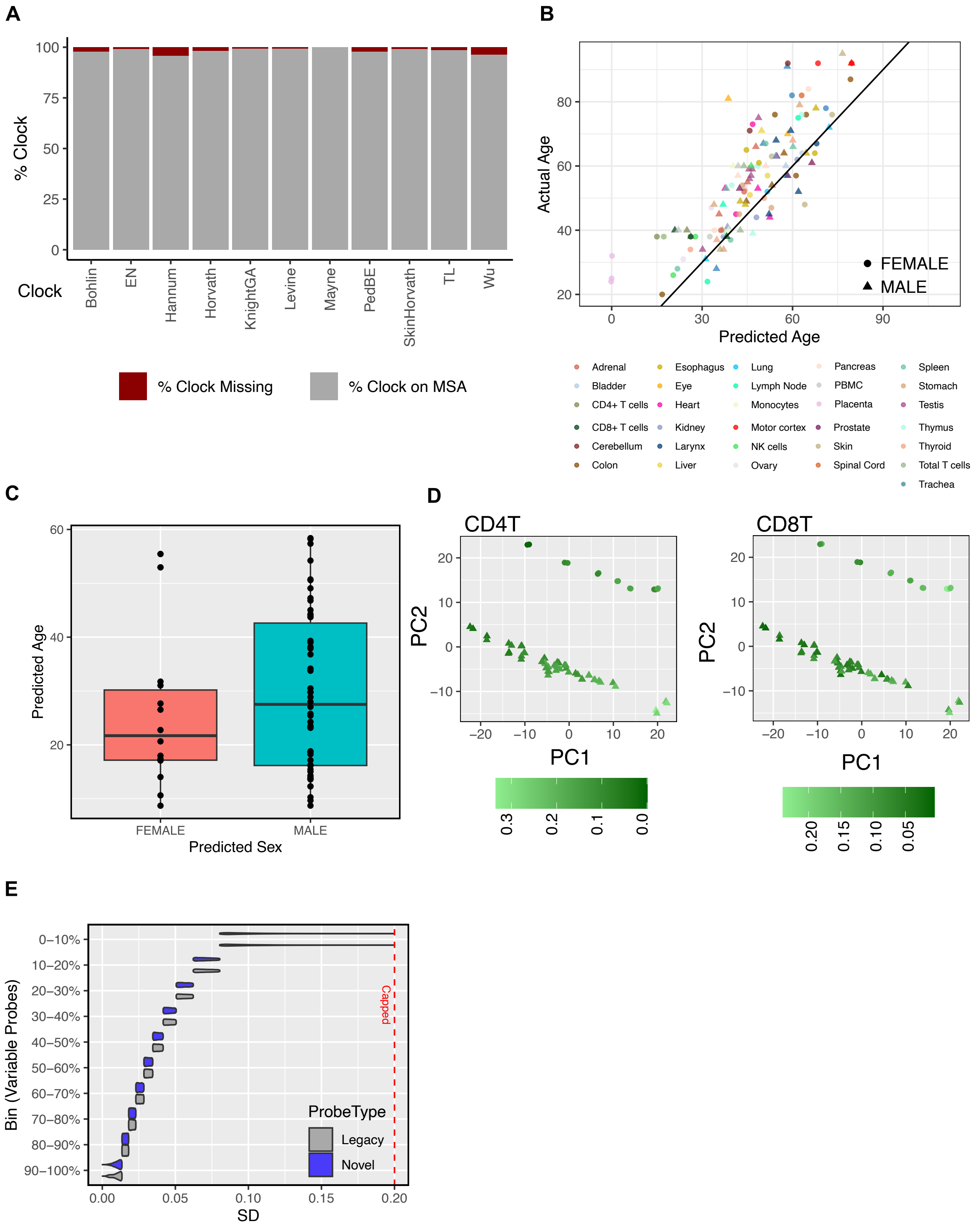


**Figure S6: Immune cell composition and interindividual whole blood methylation variation** (A) Stacked bar plots showing the percent of probes covered on MSA (grey) and the percent missing (red) for 11 epigenetic clocks (B) Horvath methylation-based age estimates compared to actual age for the tissues profiled on MSA. Age estimates are highly correlated with reported age (Pearson correlation=0.82) (C) Boxplots showing the distribution of sex and age predictions from MSA profiled whole blood samples. (B) PCA plot of whole blood methylomes with CD4T (left) or CD8T (right) cell type proportion overlaid. (B) Violin plots showing distributions of novel probe designs and reintroduced legacy probes over each decile of variably methylated probes. Novel probe designs detect similar variations.
